# A Cultured Sensorimotor Organoid Model Forms Human Neuromuscular Junctions

**DOI:** 10.1101/2021.03.03.433128

**Authors:** João D. Pereira, Daniel M. DuBreuil, Anna-Claire Devlin, Aaron Held, Yechiam Sapir, Eugene Berezovski, James Hawrot, Katherine Dorfman, Vignesh Chander, Brian J. Wainger

## Abstract

Human induced pluripotent stem cells (iPSCs) hold promise for modeling diseases in individual human genetic backgrounds and thus for developing precision medicine. Here, we generate sensorimotor organoids containing physiologically functional neuromuscular junctions (NMJs) within a cultured organoid system and apply the model to different subgroups of amyotrophic lateral sclerosis (ALS). Using a range of molecular, genomic, and physiological techniques, we identify and characterize motor neurons and skeletal muscle, along with sensory neurons, astrocytes, microglia, and vasculature. Organoid cultures derived from ALS subject iPSC lines and isogenic lines edited to harbor familial ALS mutations all show impairment at the level of the NMJ, as detected by both contraction and immunocytochemical measurements. The physiological resolution of the human NMJ synapse, combined with the generation of major cellular cohorts exerting autonomous and non-cell autonomous effects in motor and sensory diseases, may prove valuable for more comprehensive disease modeling.

## Introduction

The neuromuscular junction (NMJ) is the core synapse in the neuromuscular nervous system, with broad diagnostic and therapeutic relevance to motor neuron diseases, neuropathies, junction disorders, and myopathies. In amyotrophic lateral sclerosis (ALS), a devastating and rapidly fatal degenerative disease of the motor nervous system, degeneration of the NMJ occurs as the first pathological feature^1^. Some 90% of cases are apparently sporadic while 10% are familial and result from mutations in one of over 30 individual genes, including superoxide dismutase 1 (*SOD1*), fused-in-sarcoma (*FUS*), TAR DNA-binding protein 43 (*TARDBP*, which encodes TDP-43), profilin 1 (*PFN1),* and intronic hexanucleotide repeat expansion in the *C9orf72* gene^2^. Familial ALS genes may be broadly grouped into those that affect proteostasis, RNA binding, and axonal transport, exemplified by SOD1, TDP-43, and PFN1, respectively. The high percentage of sporadic cases, the large number of ALS genes, and the importance of human-specific splice variants and genetic backgrounds all support the need for iPSC-based modeling^3–5^.

Modeling of the NMJ using human iPSCs has proved challenging, despite multiple long-standing protocols for spinal motor neuron and skeletal muscle differentiation^6, 7^. Co-culture strategies to date have not produced NMJs from ALS subject iPSCs, although one group has successfully generated NMJs from ALS iPSC-derived motor neurons co-cultured with myoblasts differentiated from a single control iPSC line using a custom-fabricated microfluidic device^8^. Instead, most but not all reports have required the use of primary human or mouse muscle, thus losing potential genotype-specific disease contributions from the muscle^8–14^. However, the robustness and reproducibility of these models across multiple disease and control iPSC lines have not been tested, and thus their capacities for disease modeling remain unclear.

Here, we established a cultured human sensorimotor organoid model and leveraged it to probe distinct ALS variants. We performed an initial characterization of the model with five iPSC lines, obtained from two healthy controls and three ALS subjects. We found that all lines gave rise to organoid cultures containing neuronal derivatives, namely motor and sensory neurons, as well as astrocytes, and mesodermal derivatives, including vasculature, microglia, and skeletal muscle. The motor neurons and skeletal muscle connected via physiologically active NMJs, which were impaired in organoids derived from all three ALS lines. To validate the model further and pursue the ALS modeling more comprehensively, we generated isogenic pairs of iPSC lines harboring familial ALS mutations in *TARDBP*, *SOD1*, and *PFN1* and matched controls, and again found NMJ abnormalities in the organoids derived from ALS mutation-containing lines. In the gene-edited lines, we identified marked reductions in both among- and within-line variances across most metrics used to characterize the organoids. The physiological resolution of the NMJ synapse combined with the robust generation of major cellular cohorts exerting autonomous and non-cell autonomous effects in motor neuron diseases may prove valuable not only for modeling diseases but for capturing key features of their heterogeneity.

## Results

### Neuromesodermal progenitors give rise to ectodermal and mesodermal derivatives in organoid cultures

We differentiated iPSCs in suspension to form spheres, which were patterned for a week, plated at a density of 46 spheres/cm^2^, and cultured under adherent conditions for up to fifteen weeks total (Fig. 1a). Initial experiments used five iPSC lines: two controls (11a, FA0000011), two familial ALS lines (19f and MGH5b harboring C9orf72 repeat expansion and FUS mutations, respectively), and one sporadic ALS line (FA0000012) (see methods for line details). In order to generate both motor neurons and muscle, patterning was designed to produce neuromesodermal progenitors via FGF and WNT agonists as well as forskolin^15, 16^. Two day-old spheres contained neuromesodermal progenitors, identified by expression of both ectodermal (SOX2) and mesodermal (TBXT, also known as Brachyury) lineage markers (Fig. 1b)^15^. Characterization of the potency of individual spheres for generating myogenic (enriched in TBXT), neurogenic (enriched in SOX2), or neuromesodermal (equal TBXT and SOX2) lineages showed that all five lines had strong neuromesodermal potential, as measured either on the level of individual spheres (Fig. 1c) or whole-well staining (Fig. 1d). Representation of strictly myogenic or neurogenic spheres was substantially lower in all lines (Extended Data Fig. 1a,b).

**Fig. 1.**
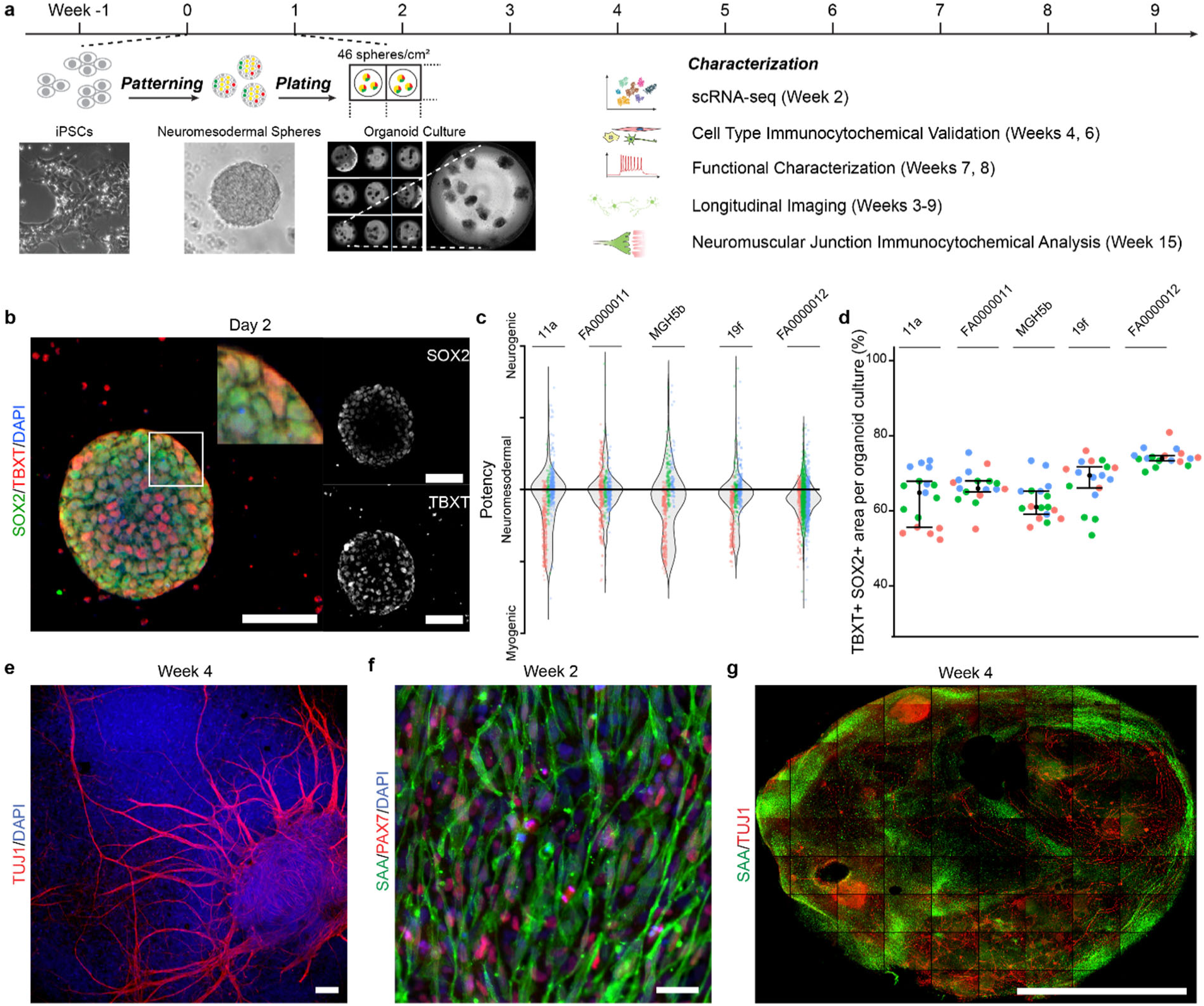
Spheres containing neuromesodermal progenitors generate neurons and myocytes. **a,** Sensorimotor organoid differentiation protocol and end-point analysis times. Spheres patterned in suspension are plated after 1 week and grown under adherent conditions for up to 15 weeks. **b,** SOX2+/TBXT+ neuromesodermal progenitors in a typical sphere, after two days in culture. Scale bar, 70 µm. **c,** Distribution of individual spheres according to expression of myogenic (TBXT), neuromesodermal (TBXT and SOX2), or neurogenic (SOX2) transcription factors, from three separate differentiations (colors). Control lines: 11a, FA0000011. ALS lines: MGH5b, 19f, and FA0000012. **d,** Quantification of the SOX2+/TBXT+ area in whole wells of the same iPSC lines. Bars indicate median and I.Q.R. (*n*=5-6 organoid cultures obtained from 3 separate differentiations in distinct colors. **e,** Ectodermal lineage cells, including TUJ1+ neurons after four weeks in culture. Scale bar, 120 µm. **f,** Mesodermal lineage cells include sarcomeric α-actinin (S.A.A.)+ myocytes, surrounded by Pax7+ cells. Scale bar, 50 µm. **g,** Whole-well confocal image of organoid cultures at four weeks showing SAA+ and TUJ1+ areas. Scale bar, 5 mm.

After the week of patterning in suspension, the number of spheres across the five lines was consistent (*n*=3 separate differentiations; one-way ANOVA, F=0.758, P=0.632, Extended Data Fig. 1c). Two to three days after plating the spheres at fixed density in matrigel-coated plates, cells migrated outward (Extended Data Video 1) to form a confluent, adherent organoid culture comprised of the plated spheres and outgrowing structures. Within the next two weeks, both ectodermal and mesodermal lineages were evident: scattered TUJ1-positive neurons, as early as day 8 (Extended Data Fig. 1d) and more densely at four weeks (Fig. 1e), as well as cells that expressed the transcription factor Pax7 in proximity to sarcomeric α-actinin-positive myocytes at two weeks (Fig. 1f and Extended Data Fig. 1e). The organoid cultures contained neurons and muscle distributed in discrete areas (Fig. 1g), and we next used single-cell sequencing of the early organoids to confirm the developing cell identities in an unbiased manner.

### Single-cell RNA-seq confirms multiple mesodermal and ectodermal cell types

We generated organoid cultures from three independent differentiations of a control iPSC line (11a) and dissociated them at two weeks, the earliest time at which we consistently observed both ectoderm and mesoderm derivatives. Using inDrop single-cell RNA-seq^17^, we sequenced libraries from 27,000 cells (9,000 cells/differentiation) at an average read depth of 35,000 reads/cell covering an average of 2,555 unique genes/cell. Cells were clustered into 25 independent groups (Fig. 2a) using the Seurat package in R^18^, and cluster identity was assessed using differentially-expressed cluster marker genes (Fig. 2b-c)^19^. Neural crest cells (clusters 0, 11, 12, 13, 14, 17, 20) were identified by expression of *FOXD3* and *SOX10*, critical transcription factors implicated in cell fate decisions of nascent neural crest cells^20^, as well as *SNAI2* and *SOX9*^21^. Cells expressing *MEST*, *MAFB*, *BMP7, SOX2,* and *OLIG3* (2, 3, 6, 8, 9, 15, 16, 18), which are important for early lineage specification of neural crest derivatives, were classified as intermediate progenitors^22^. Neuronal progenitors (5, 21) were identified by overlapping expression of intermediate progenitor markers *OLIG3*, *SOX2*, and *DLL1*^23–25^ in combination with expression of neuronal makers *MAP1B* and *TUBB3*^26^. Maturing neurons (10, 22, 24) were readily identified by strong expression of pan-neuronal markers *TUBB3* and *MAP1B*^26^. Cells in clusters 7 and 19 were classified as mesenchymal progenitors based on expression of *CXCL12*, *CD164*, *TWIST1,* and *PRRX1* (Fig. 2b-c and Supplementary Table 1)^27–29^. The clear separation of a neuronal cluster from the remaining neural crest derivatives at an early point (Fig. 2b) is consistent with single-cell RNAseq data from developing mouse neural crest showing an initial specification of a neuron fate^22^.

**Fig. 2.**
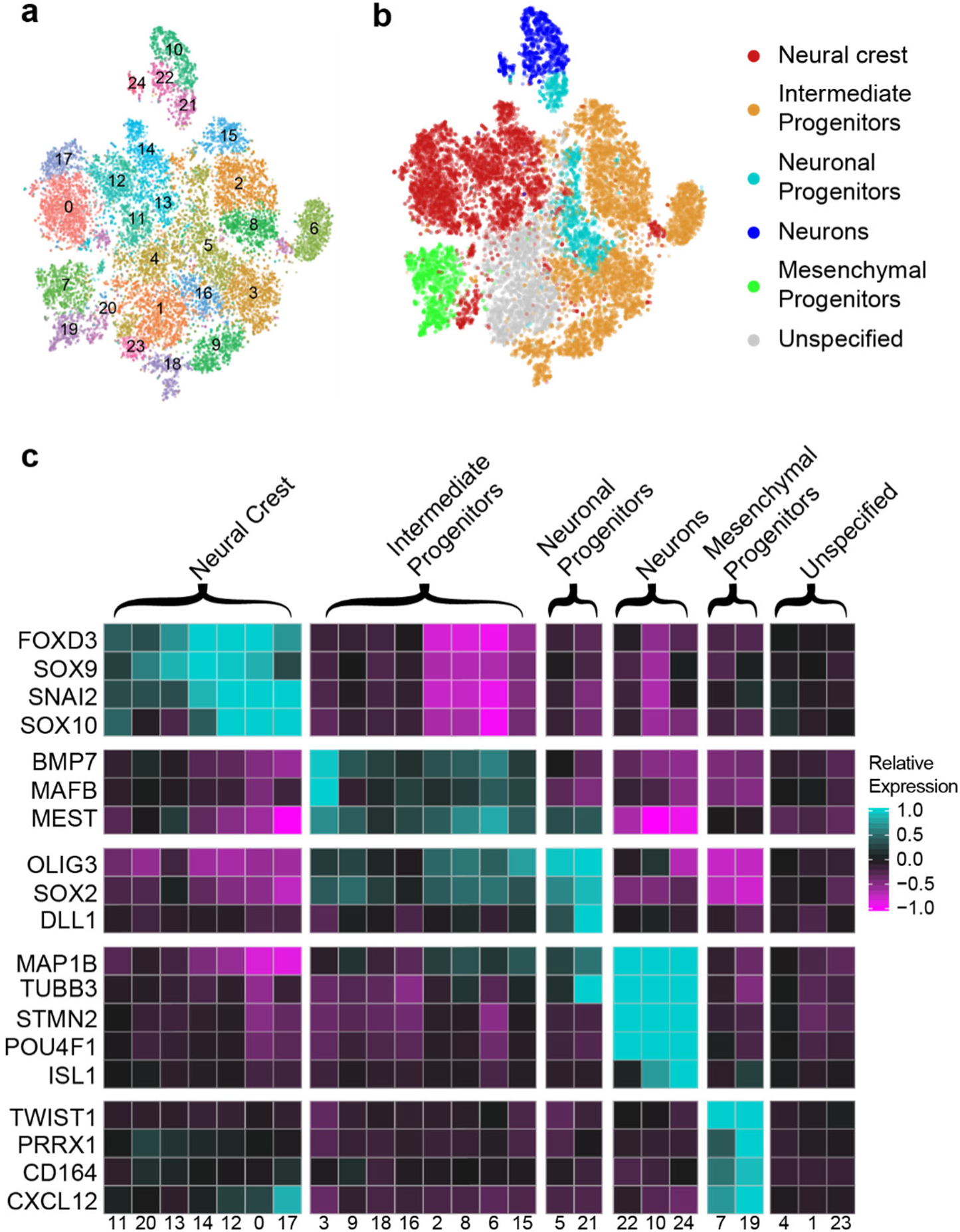
Single-cell RNA-seq analysis of early organoids confirms multiple neural and mesodermal lineages. **a,** Unsupervised cluster analysis of individual cells from early organoid cultures. Organoid cultures, three independent differentiations of the control iPSC line 11a, were dissociated at 2 weeks in culture, and 20,835 single-cell libraries were sequenced at an average read depth of 35,000 reads/cell covering an average of 2,555 unique genes/cell. Visualization shows tSNE plots of individual cells as dots and clusters of cells as colored groups. Numbers indicate cluster identity. **b,** classification of clusters into six broad cell types. Marker genes for each cell cluster were identified, and clusters were then classified into groups accordingly. Clusters that could not be unambiguously identified are labeled as unspecified. Visualization shows same tSNE plot from (**a**) recolored to reflect cell type. **c,** Expression of cluster-defining genes enriched in each cell type classification. Marker genes that are enriched in each cluster within a cell-type classification were identified. Note that the neural progenitor cell type was defined by the presence of gene subsets from both the intermediate progenitor (*OLIG3*, *SOX2*, and *DLL1*) and neuronal (*MAP1B*, *TUBB3*) cell types. Plot shows mean of scaled expression of indicated genes from within each cell cluster. High and low gene expression are indicated by cyan and magenta, respectively.

To examine effects of variable maturation state and to validate our Seurat analysis using an orthogonal approach, we used SPRING software, which infers developmental connections among cells^30^. SPRING analysis revealed 22 clusters (Extended Data Fig. 2) validated by gene ontology (GO) analysis, including neuronal (5, 8, 12), maturing skeletal muscle (7, 17, 19), vasculature development (0), and hematopoiesis (4), in general agreement with the Seurat analysis.

### Organoid cultures generate neuroectodermal derivatives including neurons and astrocytes

The single-cell RNA-seq analysis identified three distinct clusters that expressed the pan-neuronal genes *TUBB3* and *NCAM1* as well as *NEUROG1* and *NEUROG2*, which demarcate early neuronal development (Fig. 3a)^31–33^. Other expressed genes suggested specification of neuronal type: for example, ISL (*ISL1*) marks primarily motor and nociceptor neurons but also select interneuron populations^34, 35^. In motor neurons, enrichment of *STMN2* expression supports microtubule assembly for axonal outgrowth^3, 4^. In sensory neurons, the transcription factor encoding *POU4F1* (which encodes BRN3A; Seurat clusters 10 and 24) mediates essential TrkA signaling for nociceptor development^36^ and remains expressed in post-natal nociceptors (Allen Brain Atlas, http://portal.brain-map.org/). Similarly, the gene encoding for the neurofilament protein peripherin (*PRPH*) delineates unmyelinated nociceptive C-fibers and autonomic fibers in the peripheral nervous system^37^.

**Fig. 3.**
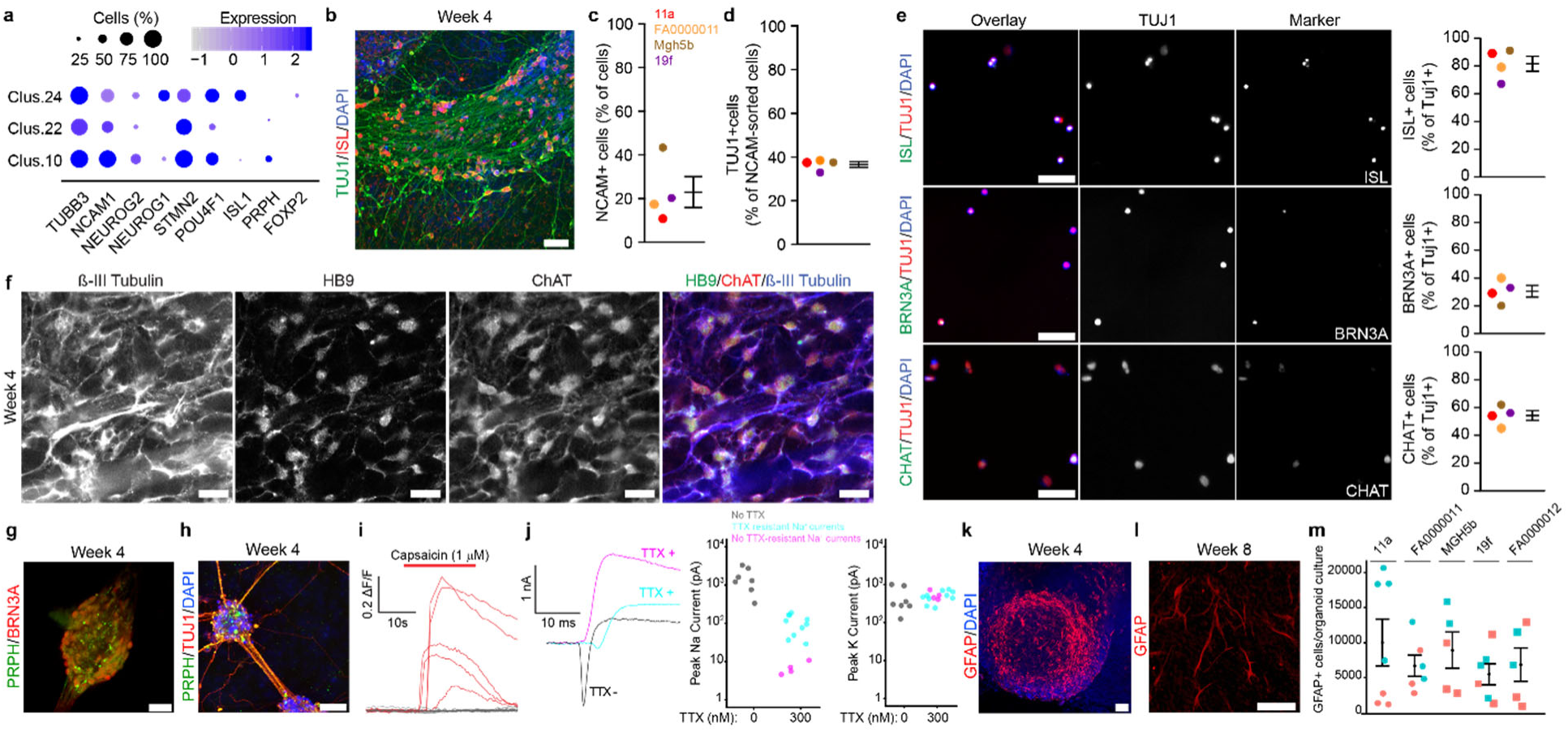
Sensorimotor organoid cultures generate cells of ectodermal lineage, including sensory and motor neurons and astrocytes. **a,** Dot plot of gene expression within neuronal clusters from single-cell RNA-seq data. Color intensity indicates expression level, and dot size indicates proportion of cells expressing each gene. **b,** Validation of ISL+/ TUJ1+ neurons in whole organoid cultures at four weeks. Scale bar, 100 µm. **c,** Quantification of NCAM+ cells after FACS purification of whole organoid cultures from four iPSC lines at four weeks. Control lines (circles): 11a (red), FA0000011 (orange). ALS lines (squares): MGH5b (brown), and 19f (purple). Bars indicate mean and S.E.M.**d,** Quantification of immunostaining for TUJ1 of FACS purified cells in (C) for four iPSC lines. Bars indicate mean and S.E.M. Control lines (circles): 11a (red), FA0000011 (orange). ALS lines (squares): MGH5b (brown), and 19f (purple) **e,** Staining (left) and quantification (right) of neuronal subtypes after FACS by expression of ISL (top), BRN3A (middle), and CHAT (bottom) from cultures of four iPSC lines. Scale bars, 50 µm. Bars indicate mean and S.E.M. Control lines (circles): 11a (red), FA0000011 (orange). ALS lines (squares): MGH5b (brown), and 19f (purple). **f,** HB9, ChAT, and β-III tubulin labeling of spinal motor neurons in four-week old organoid cultures (confirmed in 3 separate differentiations of the FA0000011 iPSC line). Scale bar, 25 µm. **g,** BRN3A and PRPH staining of ganglia within whole organoid cultures after four weeks. Scale bar, 25 µm. **h,** Ganglia connected by PRPH+/TUJ1+ axonal processes at four weeks. Scale bar, 100 µm. **i,** Sample calcium imaging traces of individual cells in ganglia-containing fields with (red) and without (grey) response to capsaicin at 6-7 weeks in culture. Capsaicin sensitivity was confirmed in three independent differentiations of two iPSC lines (11a, MGH5b). **j,** Whole cell patch clamp recordings from neurons in ganglia-containing fields at seven weeks. Example currents traces recorded in the absence (grey) or presence of 300 nM T.T.X. (cyan, cell without TTX-resistant sodium currents; magenta, cell with TTX-resistant sodium currents). Quantification of peak sodium (middle) and potassium (right) current amplitudes in the absence (grey) or presence of 300 nM T.T.X. (cyan, cell without TTX-resistant currents; magenta, cell with TTX-resistant currents). Current recordings were pooled from four separate differentiations of FA0000011 (sodium Currents: *n*=4 cells without TTX resistance, and *n*=10 cells with TTX resistance, Mann-Whitney, U=0; P=0.002; potassium currents: *n*=4 cells without TTX resistance, and *n*=10 cells with TTX resistance, U=19; P =0.945). **k,** GFAP+ astrocytes in four-week-old organoid cultures. Scale bar, 50 µm. **l,** Acquisition of typical astrocyte morphology (left) at eight weeks. Scale bar, 50 µm. **m,** Quantification of GFAP+ cells (right) in organoid cultures at six weeks. Bars indicate mean and S.E.M. (*n=*5-7 organoid cultures obtained from 2 separate differentiations identified by colors; Kruskal-Wallis, P=0.388).

To validate the single-cell RNA-seq data and the expression of ISL in TUJ1-positive neurons of more mature organoids, we first immunostained whole-organoid cultures after four weeks and found groups of ISL-positive neurons (Fig. 3b). To assess multiple subpopulations within the same organoid culture, we then used a two-step process of fluorescence-activated cell sorting (FACS) followed by immunocytochemistry immediately after plating the purified cells to quantify neuronal subtypes. We found that NCAM-based FACS selected 23.0 ± 7.1% (mean ± S.E.M.) of cells (Fig. 3c and Extended Data Fig. 3), and subsequent staining for TUJ1 (Fig. 3d) revealed that 38.0 ± 1.3% of the NCAM-purified cells (mean ± S.E.M., Fig. 3d) were neuronal. NCAM, which has been useful for prior FACS-based neuronal purification^3^, is also expressed in some non-neuronal cells – including embryonic skeletal muscle^38^, consistent with neuronal and non-neuronal NCAM expression in our single-cell RNA-seq data (Fig. 3a; Extended Data Dataset 1 for Fig. 2). Combining the FACS purification and staining, 8.7 ± 2.7% of total cells were NCAM- and TUJ1-positive neurons (mean ± S.E.M.), in agreement with the 8.5% of cells in neuronal clusters identified by single-cell RNA-seq (Supplementary Table 1).

In the NCAM-purified and TUJ1-positive neurons, we identified a large percentage of ISL-positive cells (81.5 ± 5.5%, mean ± S.E.M.) (Fig. 3e). To distinguish among ISL-expressing neuronal subtypes, we stained and quantified BRN3A-positive nociceptors (30.5 ± 4.2%, mean ± S.E.M.) and choline O-acetyltransferase (ChAT)-positive motor neurons (54.3 ± 3.5%, mean ± S.E.M.) (Fig. 3e). The cumulative proportion of nociceptor and motor neurons (86%) was concordant with the overall expression of ISL (81.5 ± 5.5%, mean ± S.E.M., Fig. 3e) and consistent with the specificity of ISL for these two neuronal subtypes. Additionally, labeling of whole organoid cultures at four weeks confirmed the presence of ChAT+ neurons expressing the motor neuron marker HB9 (*MNX1*, also known as HoxB9) (Fig. 3f, Extended Data Fig. 4), an established profile demarcating spinal motor neurons^39^. Quantitative RT-PCR at ten weeks showed persistent expression of nociceptor (*POU4F1* and *PRPH*, Extended Data Fig. 5a) as well as motor neuron (*LHX3*, *CHAT*, Extended Data Fig. 5b) markers.

**Fig. 4.**
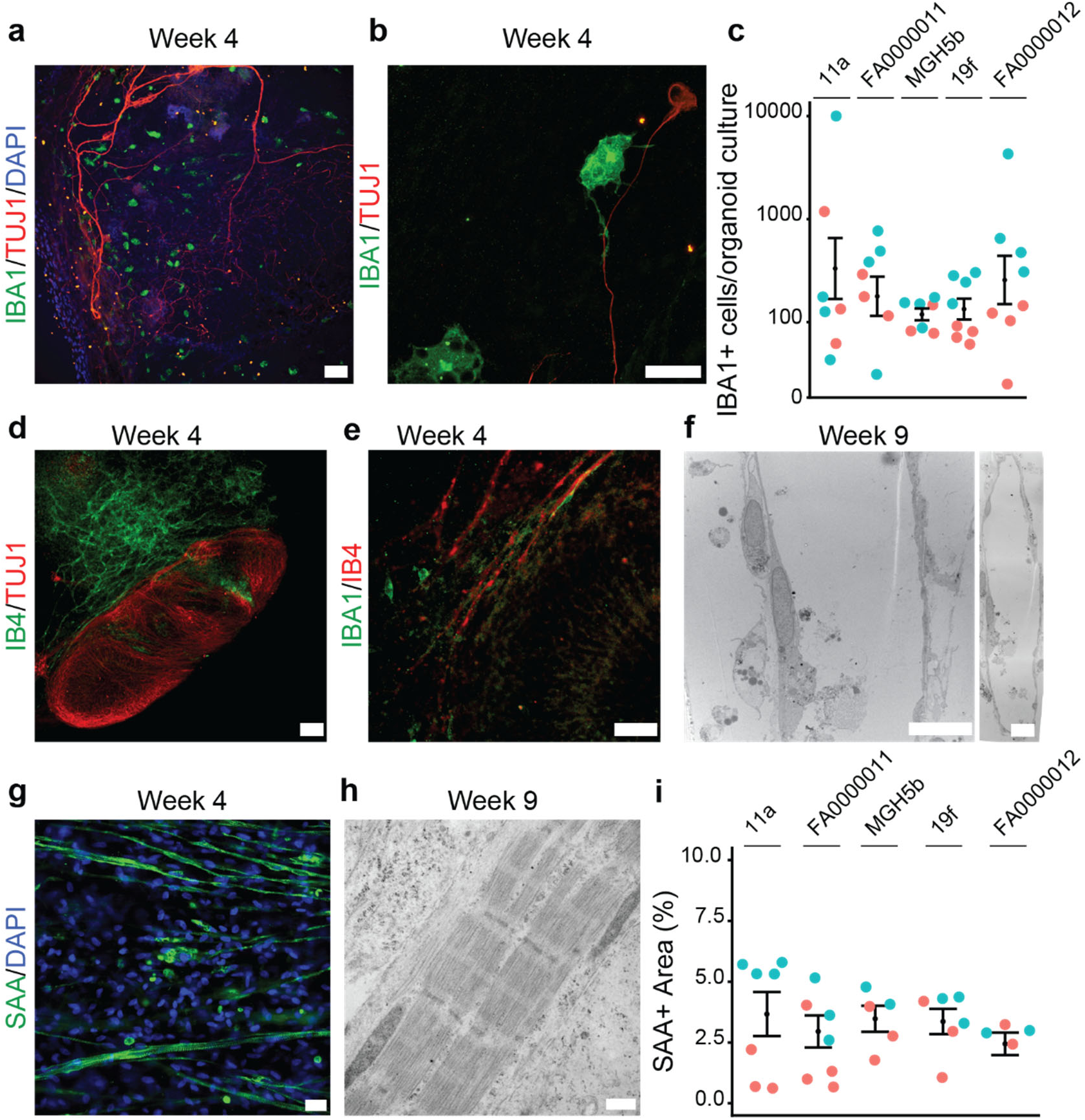
Organoid cultures differentiate into mesodermal lineage cells that include IBA1-expressing microglia, vasculature, and skeletal muscle. **a,** Microglia in sensorimotor organoids immunolabeled with IBA1 at four weeks. Scale bar, 100 µm. **b,** High magnification image of an IBA1+ microglial cell and TUJ1+ axon at four weeks of culture. Scale bar, 50 µm. **c,** Quantification of IBA1+ cells in organoid cultures at six weeks. Bars indicate mean and S.E.M (*n=*6-8 organoid cultures obtained from 2 separate differentiations identified by colors; Kruskal-Wallis, P=0.613). **d,** Microvasculature stained with IB4 nearby TUJ1+ neurons at four weeks in culture. Scale bar, 100 µm. **e,** IB4+ microvasculature adjacent to IBA1+ microglia at four weeks in culture. Scale bar, 50 µm. **f,** Electron micrograph of endothelial-lined microvessels at high (left) and low (right) magnification after nine weeks in culture. Scale bars, 10 µm. **g,** Elongated and striated SAA+ myotubes containing peripheral nuclei at four weeks. Scale bar, 100 µm. **h,** Electron micrographs of skeletal muscle at nine weeks showing sarcomeric organization with distinct Z lines, A and I bands, and M bands within the H zone. Scale bar, 500nm. **i,** Quantification of percent of SAA+ muscle area in organoid cultures at six weeks. Bars indicate mean and S.E.M. (*n*=5-6 organoid cultures obtained from 2 separate differentiations identified by colors; Kruskal-Wallis, P=0.642). Control lines: 11a, FA0000011. ALS lines: MGH5b, 19f, and FA0000012.

**Fig. 5.**
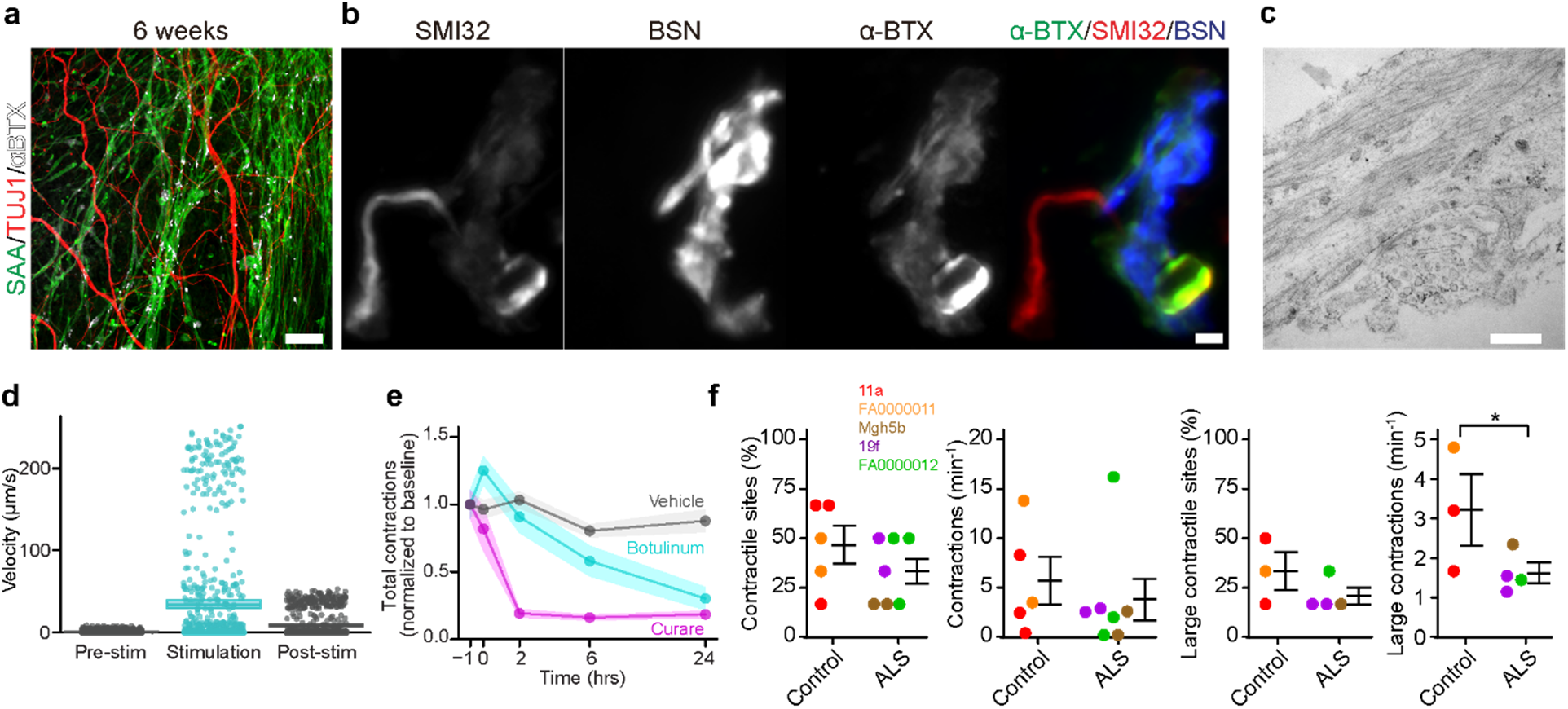
Sensorimotor organoids form functional NMJs, which are impaired in ALS cultures. **a,** TUJ1+ neurons juxtaposed with SAA+ and α-bungarotoxin (αBTX)+ skeletal muscle at six weeks in culture. Scale bar, 100 µm. **b,** NMJs at high magnification depicting SMI32+ neuronal terminals and presynaptic B.S.N. staining that abuts postsynaptic αBTX staining. Scale bar, 2 µm. **c,** Electron micrograph of an NMJ showing synaptic densities that separate presynaptic vesicles (bottom right) from longitudinally cut muscle fibers (top) at nine weeks in culture. Scale bar, 500nm. **d,** Quantification of muscle contraction triggered by optogenetic activation of neurons. Neurons expressing *hSYN*::ChR2 evoke rapid movement of muscle fibers when stimulated with blue light. **e,** Pharmacology of muscle contractions in sensorimotor organoids. Spontaneous muscle contractions in eight-week-old cultures are inhibited by curare and botulinum toxin, the latter indicating dependence on synaptic function. Inhibition of contractions by each drug was assessed in five independent differentiations of four iPSC lines (11a, MGH5b, 19f, and FA0000012). Plots show mean and 95% confidence intervals as line and shaded area, respectively. **f,** Unbiased analysis of contraction site and frequency. Skeletal muscle contractions at six sites per organoid culture were counted and pooled from three independent differentiations of five iPSC lines at 7-8 weeks. Percentage of contractile sites and contraction frequency per organoid culture were quantified separately for all contractions (left two panels, *n*=5 control and 7 ALS organoids, ANOVA type 3, F=1.458, P=0.255), and large contractions (right two panels, *n*=3 control and 4 ALS organoid cultures, ANOVA type 3, P=0.035). Circles and squares indicate control and ALS iPSC lines, respectively: 11a (red), FA0000011 (orange), MGH5b (brown), 19f (purple), and FA000012 (green).

Based on the expression of the sensory neuron markers, we looked for structures in the organoid cultures that contained sensory neurons and resembled dorsal root ganglia^40^. BRN3A-positive sensory neurons formed ganglia-like clusters (Fig. 3g) expressing PRPH-positive neuronal processes (Fig. 3h), resembling their *bona fide* counterparts; in contrast, neurons outside the ganglia did not express PRPH. We then investigated whether the candidate sensory neurons expressed functional nociceptor-specific channels and ionotropic receptors that serve key roles in pain sensory transduction. We first tested for the functional expression of TrpV1 receptors using capsaicin, the pungent ingredient in hot chili peppers^41^. Using calcium imaging of fields containing ganglia-like structures, bath application of capsaicin elicited robust calcium flux in 169 cells pooled from three independent organoid differentiations (Fig. 3i). We next assessed the presence of tetrodotoxin-resistant (TTX-R) sodium currents, mediated by Na_V_1.8 and Na_V_1.9 nociceptor-specific ion channels^42, 43^. We recorded sodium currents from ganglia in the absence or presence of 300nM TTX using whole-cell patch clamp (Fig. 3j). In addition to robust sodium currents in the absence of TTX (1300 pA [982-2165] median peak sodium amplitude [I.Q.R.]; *n*=7 cells), we observed TTX-R currents with slower activation kinetics, as observed in heterologous studies of Na_V_1.8 channels^42^, in a large proportion of cells (TTX-R-containing cells: 105 pA [55-153], *n=*10/14 cells; TTX-R-lacking cells: 6 pA [5-8], *n=*4/14 cells; median peak sodium amplitude [I.Q.R.]; Mann-Whitney test U=0, two-tailed, P=0.002). We observed no difference in voltage-gated potassium currents in the same cells (TTX-R lacking cells: 303 [277-646] pA, *n*=7 cells; TTX-R-containing cells: 492 [429-613] pA, *n=*10/14 cells; TTX-R-lacking cells: 484 pA [438-572], *n=*4/14 cells; median steady-state potassium amplitude [I.Q.R.]; Mann-Whitney test U=19, two-tailed, P=0.945).

Consistent with the overlapping developmental pathways for neurons and astrocytes, as well as prior studies showing astrocytes in brain organoids^44^, we anticipated finding astrocytes in the sensorimotor organoids. Indeed, we identified GFAP-positive cells by immunocytochemistry as early as four weeks (Fig. 3k-m). Early GFAP-positive cells migrated out of the spherical structures but did not show an obvious astrocyte morphology until eight weeks in culture (Fig. 3l and Extended Data Fig. 5c), and quantification of GFAP-positive staining from whole organoid cultures by confocal imaging showed no significant differences among the lines (*n*=5-7 organoid cultures obtained from 2 separate differentiations; Kruskal-Wallis, P=0.389; Fig. 3m and Extended Data Fig. 5c).

### Organoid cultures yield mesodermal derivatives including microglia, endothelial cells, and skeletal muscle

During embryonic development, microglia originate from the extraembryonic yolk sac, migrate to the developing spinal cord, and remain after the formation of the blood-brain barrier^45, 46^. The mesodermal origin of microglia, together with hematopoietic clusters in the single-cell RNA-seq data, led us to test for the presence of microglia in organoid cultures. We observed IBA1-positive cells bearing a microglia-like morphology (Fig. 4a) close to TUJ1-positive neurons (Fig. 4b) after four weeks of culture. Quantification of the primitive microglial population in cultures from five pluripotent stem cell lines (Fig. 4c and Extended Data Fig. 6a) at six weeks showed no significant difference among the iPSC lines (*n*=6-8 organoid cultures obtained from 2 separate differentiations; Kruskal-Wallis, P=0.613; Fig. 4c and Extended Data Fig. 6a). Because IBA1 and other individual markers for microglia are expressed in multiple myeloid derivatives, we used qPCR at ten weeks in culture to confirm the persistent expression of additional microglial markers, including *TMEM119* and the fractalkine receptor *CX3CR1* (Extended Data Fig. 6b)^47, 48^.

**Fig. 6.**
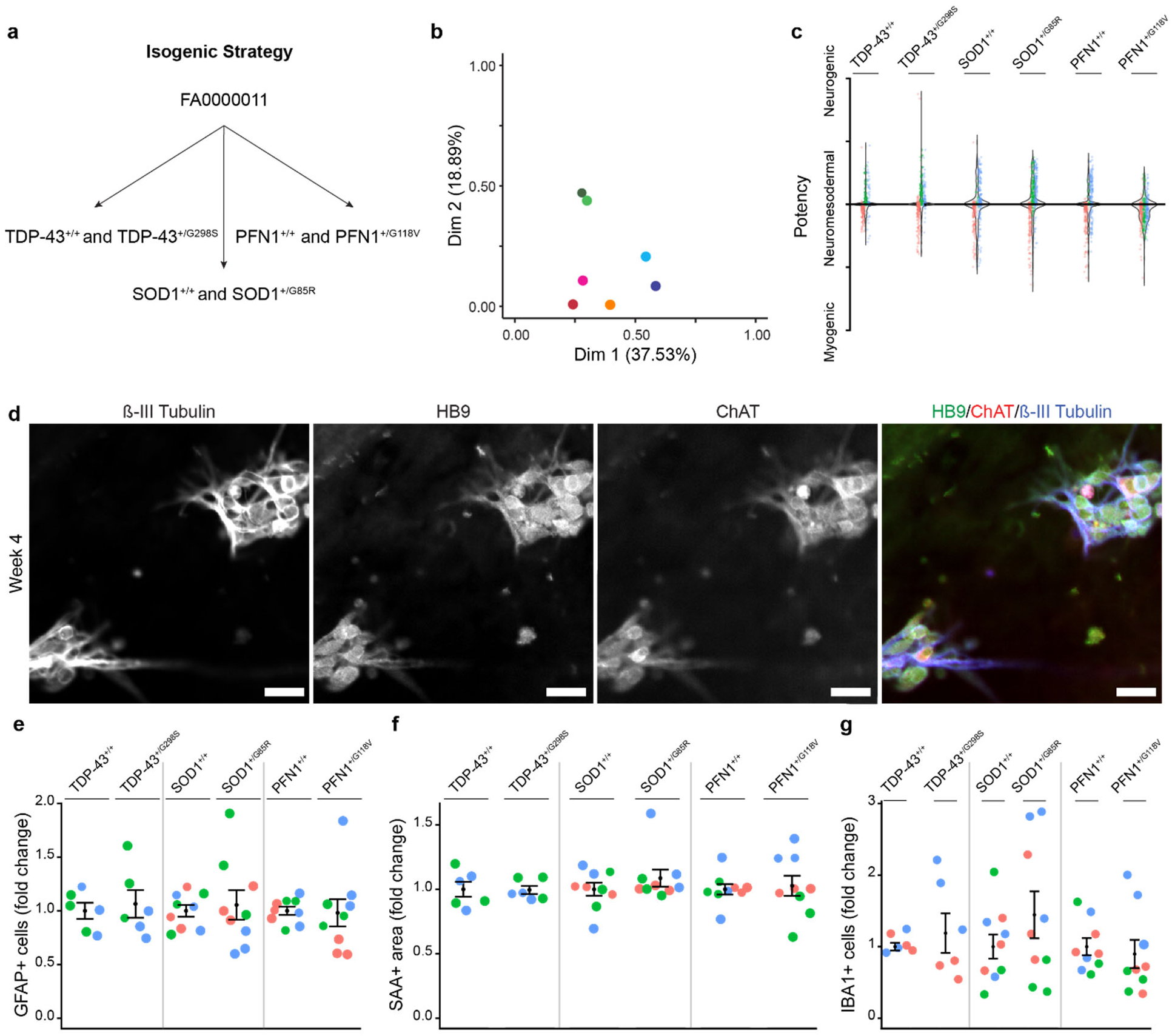
Gene-edited iPSC lines are paired with their isogenic control and generate sensorimotor organoids with similar composition. **a,** Strategy for CRISPR/Cas9-mediated insertion of point mutations in either *TARDBP* (TDP-43), *SOD1*, or *PFN1* into the FA0000011 control line. **b,** Multiple correspondence analysis of TDP-43^+/+^ (dark green), TDP-43^+/G298S^ (light green), SOD1^+/+^(dark blue), SOD1^+/G85R^ (light blue), PFN1^+/+^ (red), PFN1^+/G118V^ (pink), and the original healthy control line FA0000011 (orange). **c,** Distribution of individual spheres according to their expression of myogenic (TBXT) or neurogenic (SOX2) transcription factors from three separate differentiations (colors). **d,** Spinal motor neurons expressing HB9, ChAT, and β-III Tubulin, in organoid cultures of the SOD1^+/G85R^ iPSC line at four weeks (confirmed in three separate differentiations). **e,** Quantification of GFAP+ cells (right) in organoid cultures at six weeks, normalized to paired isogenic control, per differentiation. Bars indicate mean and S.E.M. (*n*=6-9 organoid cultures obtained from 2-3 separate differentiations identified by colors; Kruskal-Wallis, P=0.934). **f,** Quantification of percent of SAA+ area (right) in organoid cultures at six weeks, normalized to paired isogenic control, per differentiation. Bars indicate mean and S.E.M. (*n*=6-9 organoid cultures obtained from 2-3 separate differentiations identified by colors; Kruskal-Wallis, P=0.432). Bars indicate mean and S.E.M. **g,** Quantification of IBA1+ cells (right) in organoid cultures at six weeks. Bars indicate mean and S.E.M. (*n*=6-9 organoid cultures obtained from 2-3 separate differentiations identified by colors; Kruskal-Wallis, P=0.983).

The presence of microglia suggested the existence of primary vasculature, as the two share a common embryonic origin^45^. Staining with tomato isolectin B4 (IB4), which labels endothelial cells in microvasculature^49^, showed a characteristic reticulated pattern in proximity to neurons (Fig. 4d) and in some cases coursing adjacent to IBA1-positive microglia (Fig. 4e). Although IB4 also marks non-peptidergic nociceptor neurons^40^, the observed labeling pattern was not typical for neurons. To confirm the presence of vasculature, we detected expression of the vascular endothelial cadherin *CDH5* by qPCR at ten weeks (Extended Data Fig. 6b), and electron microscopy revealed the paradigmatic feature of blood vessels, namely endothelial cells lining a lumen (Fig. 4f).

Finally, given the presence of myocytes early in organoid culture (Fig. 1e), as well as the myogenic clusters in the RNA-seq data, we asked whether further muscle development and maturation occurred. We observed elongated, thin, and fused striated cells with peripheral nuclei, indicative of myotube maturity in skeletal muscle^50^ (Fig. 4g). Electron microscopy highlighted the archetypal ultrastructural features of skeletal muscle, including Z-lines within I bands and H-bands with M-lines (Fig. 4h)^51^. The total area of skeletal muscle occupied a consistent percentage of the total well area (13×10^7^ µm^2^) at six weeks in culture (*n*=5-6 organoid cultures from 2 separate differentiations; Kruskal-Wallis, P=0.642; Fig. 4i and Extended Data Fig. 6c). Expression of key skeletal muscle regulators *MYF5* and *MYOG*^16^ was also detected by qPCR at ten weeks in culture (Extended Data Fig. 6d).

### NMJs produce neuron-dependent skeletal muscle contractions, which are reduced in ALS-derived organoid cultures

To assess the structural and functional integration of motor neurons and skeletal muscle, we looked for NMJs, the critical synapse in neuromuscular disease. The organoids generated a complex network of neurons and muscle punctated by α-bungarotoxin (α-BTX)-binding nicotinic acetylcholine receptors (Fig. 5a), which can form clusters before muscle innervation but are reorganized by synaptic initiation and activity at the NMJ^52^. We observed clusters of postsynaptic α-BTX staining opposite the presynaptic neuronal marker bassoon (BSN, Fig. 5b)^53^, and electron microscopy confirmed the presence of synaptic densities flanked by presynaptic vesicles and longitudinally-cut postsynaptic muscle fibers (Fig. 5c)^50^.

Beginning at eight weeks in culture, we observed the onset of spontaneous muscle contractions (Extended Data Video 2), as expected given the incipient motor neuron network activity in both purified primary rodent and purified human stem cell-derived motor neurons^54^ as well as the spontaneous transmitter release from motor neurons yielding muscle contraction during *in vivo* development^55^. Contractions persisted for up to a month in culture and occurred consistently in all lines studied. To confirm that motor neuron activity was sufficient to elicit contractions, we expressed the blue-shifted channelrhodopsin TsChR2 under the neuronal-specific *SYN1* promoter^56^, and illumination elicited immediate and robust muscle contraction (Fig. 5d and Extended Data Video 3).

The pharmacology of the mammalian NMJ is well-characterized and serves as the basis for paralytics used broadly in clinical anesthesia^57^. To assess whether motor neuron signaling was necessary for the spontaneous muscle contractions, we applied the presynaptic and postsynaptic blockers botulinum toxin and curare, respectively, to organoid cultures. We selected individual sites that exhibited large, spontaneous muscle contractions from the five iPSC lines, including control and ALS lines, at 8-9 weeks in culture and quantified contractions using the Fiji (ImageJ 1.52p) optic flow plug-in^58^ to quantify pixel movement associated with each contraction (Extended Data Video 4). The application of curare, an antagonist of postsynaptic skeletal muscle nicotinic acetylcholine receptors, quickly abolished muscle contractions (Fig. 5e and Extended Data Fig. 7a). To verify that the spontaneous muscle contractions resulted from presynaptic motor neuron activation, we showed that botulinum toxin also blocked contractions (Fig. 5e and Extended Data Fig. 7b)^59^, albeit after a longer delay as occurs clinically^60^.

**Fig. 7.**
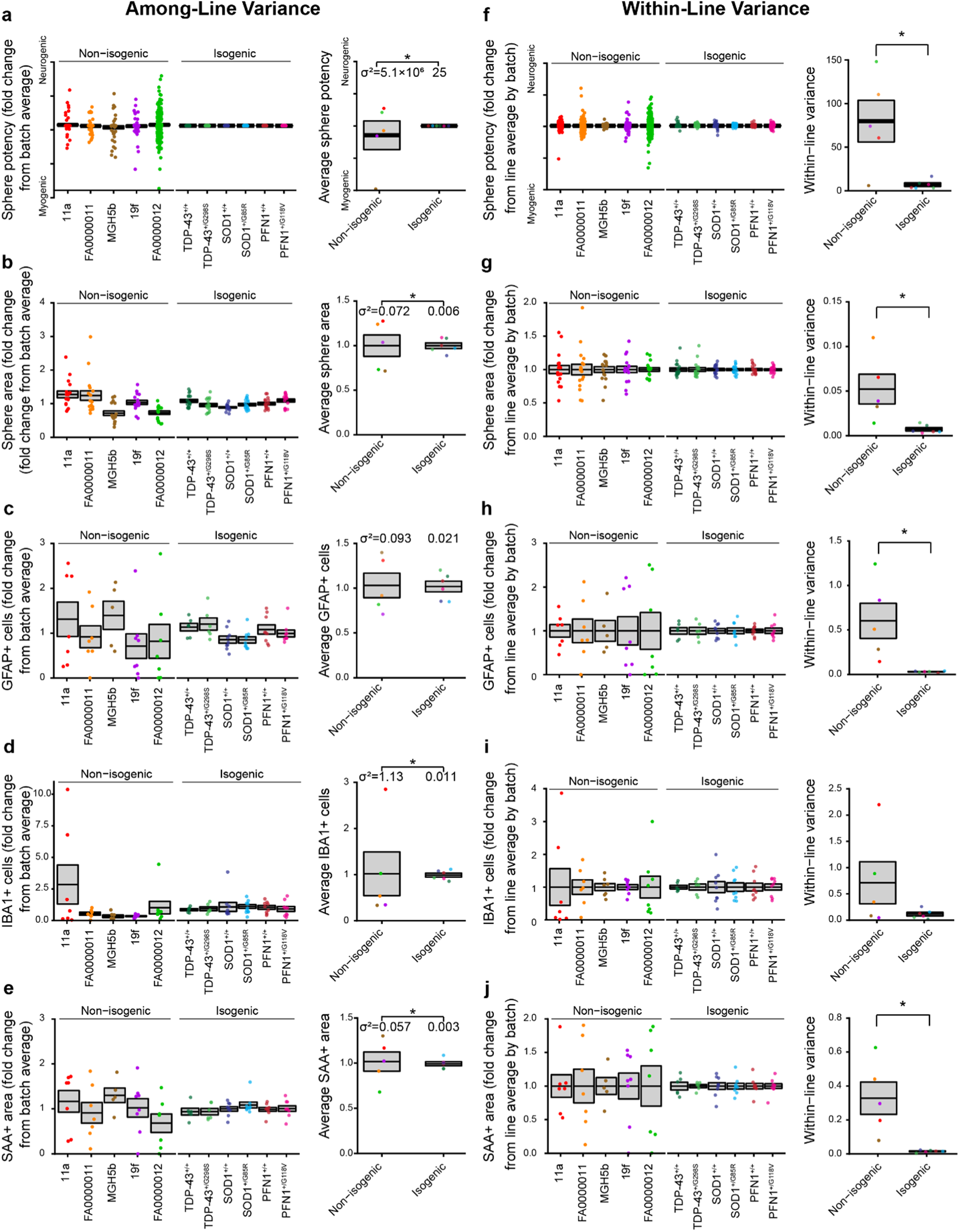
Isogenic iPSC lines reduce among- and within-line variability of organoid cultures. Analysis of among-line (**a-e**) and within-line (**f-j**) variance for individual sphere potency (**a,f**) and area (**b,g**), as well as GFAP+ cells (**c,h**), IBA1+ cells (**d,i**), and SAA+ area (**e,j**) within mature organoid cultures from 2-3 independent differentiations. **a-e**, l*eft*, Mean and SEM of fold change from batch average, along with points for individual spheres (a,b) or cultures (c-e). *Right*, Mean and SEM among non-isogenic and isogenic lines with points for each unique iPSC line. Significance assessed by F-test for Equality of Variances, with group variances (σ^2^) indicated at bottom, for *n*=5 non-isogenic, 6 isogenic lines: **a**, F=2.0×10^5^, P=6.8×10^-13^); **b**, F=2.0×10^5^, P=6.8×10^-13^; **c**, F=4.37, P=0.137; **d**, F=106.33, P=1.0×10^-4^; **e**, F=18.90, P=0.006. **f-j**, l*eft*, Mean and SEM of fold change from batch average within-line, along with points for individual spheres (f,g) or cultures (h-j). *Right*, Mean and SEM of within-line variances for non-isogenic and isogenic lines with points for each unique iPSC line. Significance assessed by one-way ANOVA for *n*=5 non-isogenic, 6 isogenic lines: **f**, (non-isogenic vs isogenic) 79.9 ± 24.0 vs 7.0 ± 2.1, F=11.21, P=0.009; **g**, 0.052 ± 0.017vs 0.007 ± 0.002, F=8.825, P=0.016; **h**, 0.603 ± 0.198 vs 0.030 ± 0.004, F=10.25, P=0.011; **i**, 0.711 ± 0.401 vs 0.114 ± 0.035, F=2.70, P=0.135; **j**, 0.328 ± 0.095 vs 0.015± 0.003, F=13.2, P=0.005. Colors indicate individual iPSC lines.

Although sites with large and frequent contractions could be found in cultures from both ALS and control lines, we subjectively noted fewer contractions in organoid cultures derived from ALS iPSC lines. This observation led us to hypothesize that the organoids captured an impairment of NMJ function in ALS lines, as loss of NMJs is one of the earliest hallmarks of ALS^1^. To evaluate the suggested reduction of contractions in ALS organoids, we performed an unbiased contractility assay in 7-8-week-old organoid cultures. While analysis sites for the pharmacology assay were specifically selected for high contraction rates, sites for the contractility assay were selected based only on the presence of muscle fibers detectable in brightfield and blind to line identity of the two control and three ALS iPSC lines using six individual sites per organoid culture from three independent differentiations of each line. We quantified the rate of skeletal muscle contraction across the control and ALS iPSC lines (Fig. 5f) by recording brightfield videos of muscle, identifying fluctuations in pixel intensity using automated frame subtraction, and then manually verifying whether fluctuations corresponded to muscle contraction all blind to line identity. We detected contractions in 12 out of the 15 organoid cultures analyzed (five out of six of the control and seven out of nine of the ALS organoid cultures). The percentage of contractile sites per organoid culture, as well as the overall contraction rate, were similar across groups (*n*=5 control and 7 ALS organoid cultures obtained from 3 separate differentiations, Percentage of contractile sites: ANOVA type 3, F=1.458, P=0.255; Contraction rate: ANOVA type 3, F=0.343, P=0.571). We observed large contractions (encompassing over half a field of view) in seven out of 15 organoid cultures (three out of six of the control and four out of nine of the ALS organoid cultures). Distinguishing between small and large contractions revealed that, despite the similar percentages of sites with large contractions (*n*=3 control and 4 ALS organoid cultures obtained from 3 separate differentiations, ANOVA type 3, F=1.753, P=0.243), there was a reduction of 72.8% in the frequency of large contractions in ALS compared to control cultures (*n*=3 control and 4 ALS organoid cultures obtained from 3 separate differentiations, ANOVA type 3, F=8.289, P=0.035).

### Gene editing of iPSC lines reduces within- and among-line variability of sensorimotor organoids

While the five iPSC lines consistently generated organoid cultures containing ALS-relevant cell types, namely, motor neurons, muscle, astrocytes, and microglia, we were concerned that the large within- and among-line variation observed in cell counts could decrease sensitivity or confound the interpretation of disease modeling studies. To reduce the influence of individual genetic backgrounds on the observed variability of sensorimotor organoids, we designed a strategy of editing three familial ALS mutations into a single control line, yielding three isogenic pairs of ALS and matched control lines. We used CRISPR/Cas9 to introduce the point mutations *TARDBP^G298S^*, *SOD1^G85R^*, and *PFN1^G118V^* into the FA0000011 control line. Individual clones with a single heterozygous mutation and no modification of the opposite allele were selected and expanded to generate the following iPSC lines: TDP-43^+/G298S^, SOD1^+/G85R^, PFN1^+/G118V^, whereas clones with no detectable editing were selected as paired controls (TDP-43*^+/+^*, SOD1*^+/+^*, PFN1*^+/+^*) (Fig. 6a). Whole exome sequencing of the isogenic ALS lines and paired controls detected variants, but none predicted by off-target effects of the guide RNAs (guide RNAs, as well as predicted off-targets, are in Supplementary Table 2; a full list of variants predicted to affect gene expression is available in Supplementary Dataset 1)^61^. Multiple correspondence analysis and hierarchical clustering of the lines using exome sequencing variants demonstrated that isogenic pairs clustered together compared to unpaired lines (Fig. 6b; Extended Data Fig. 8).

**Fig. 8.**
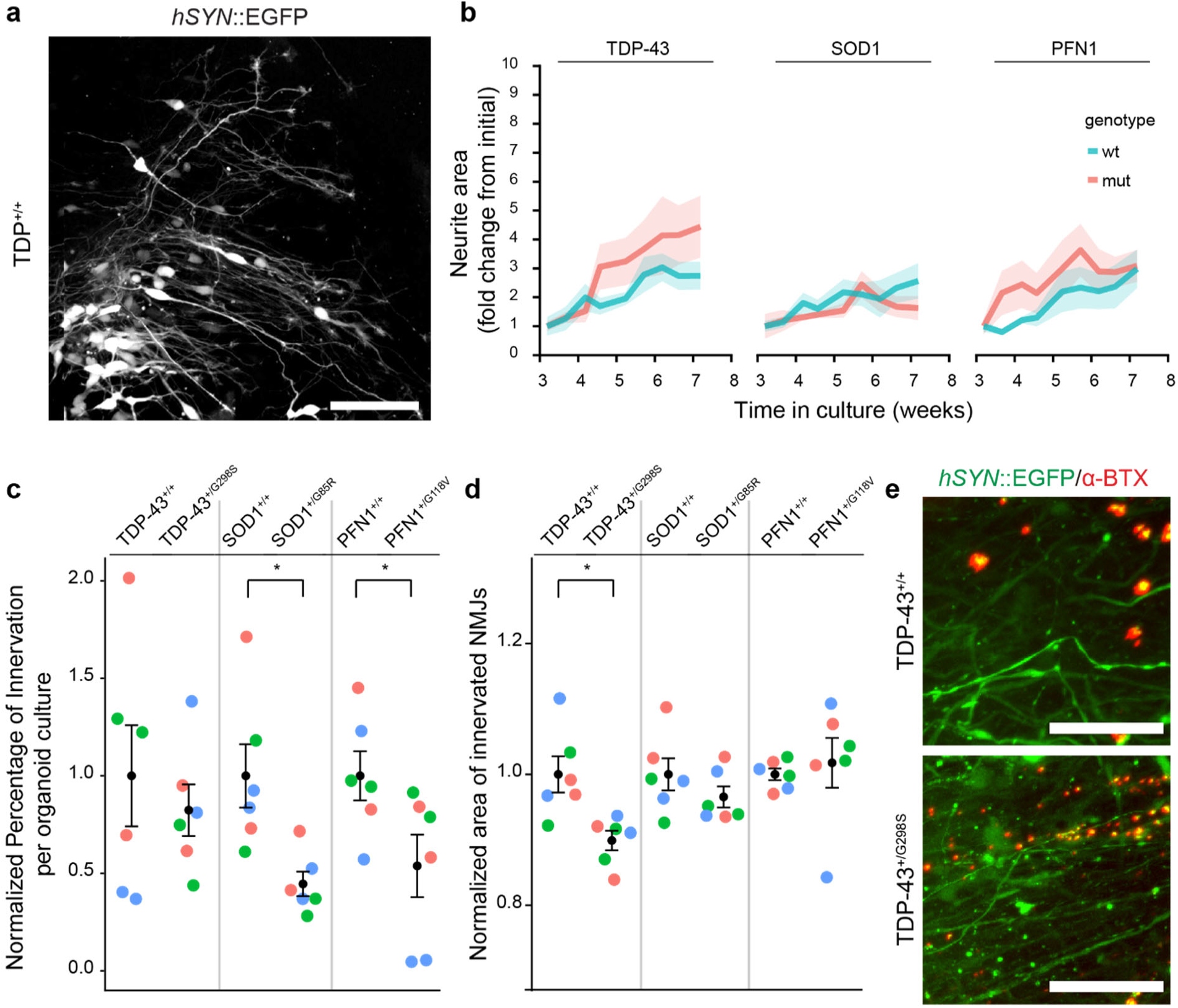
Isogenic ALS mutations show distinct NMJ phenotypes. **a,** Live neuronal labeling by expression of *hSYN*::EGFP delivered by an AAV9 vector. Scale bar, 100 µm. **b,** Quantification of neurite area as fold change of initial, from three to nine weeks in culture. Line indicates mean and shaded area indicates S.E.M.(effect of time on neurite area, *n*=6 organoid cultures obtained from 3 separate differentiations, linear model, F=8.743, P=1.3×10^-10^, individual comparisons: TDP-43 pair: F=37.040, P=2.0×10^-8^; SOD1 pair: F=16.3725, P=1.0×10^-4^; PFN1 pair: F=18.639, P=3.6×10^-5^; interaction between time and iPSC line on neurite area, *n*=6 organoid cultures obtained from 3 separate differentiations, linear model, F=0.562, P=0.985). **c,** Normalized percentage of NMJ innervation at three months in culture (normalized to the paired isogenic controls). Bars indicates mean and S.E.M. (*n*=6 organoid cultures obtained from 3 separate differentiations identified by colors, Univariate ANOVA: F=5.938, P=6.3×10^-4^; separate two-tailed t-test; TDP-43 pair: P=0.560; SOD1 pair: P=0.010; PFN1 pair: P=0.047). **d,** Normalized area of the innervated NMJs at three months in culture (normalized to the paired isogenic controls). Bars indicate mean and S.E.M. (*n*=6 organoid cultures obtained from 3 separate differentiations identified by colors, Univariate ANOVA: F=3.001, P=0.026; separate two-tailed t-test; TDP-43 pair: P=0.009; SOD1 pair: P=0.203; PFN1: P=0.659). **e,** Example of the reduction in NMJ area observed in the TDP-43^+/G298S^ iPSC line compared to the isogenic control. Scale bar, 100 µm.

To test the capacity of the six isogenic lines to make sensorimotor organoids, we first profiled the individual spheres from multiple independent differentiations for neuromesodermal potency, as for the initial five iPSC lines (Fig. 1d), and found that all lines generated large numbers of neuromesodermal spheres. All lines had the potency to generate both myogenic and neurogenic lineages, and predominantly made multipotent spheres (Fig. 6c). Similarly to the non-isogenic lines, the six isogenic lines all generated ChAT+, HB9+ spinal motor neurons (Fig 6d; Extended Data Fig. 9). Quantification of cell types at six weeks of organoid culture showed no differences in GFAP+ cells (*n*=6-9 organoid cultures obtained from 2-3 separate differentiations; Kruskal-Wallis, P=0.934; Fig. 6e), SAA+ area (*n*=6-9 organoid cultures obtained from 2-3 separate differentiations; Kruskal-Wallis, P=0.432; Fig. 6f), and IBA1+ cells (*n*=6-9 organoid cultures obtained from 2-3 separate differentiations; Kruskal-Wallis, P=0.983; Fig. 6g). Thus, the culture protocol reliably generated a consistent number of several different cell types from the isogenic paired lines.

Comparing the six isogenic lines to the original five, we noticed reduced variability in many of the early and late metrics. We examined the variation by plotting different sphere and cell type measurements for each line and quantified the within-line and among-line variance components. For the individual spheres, we analyzed the sphere neuromesodermal potency and area after two days of culture; for the organoid cultures, we analyzed astrocyte, microglia, and muscle generation by immunocytochemistry after six weeks in culture. The among-line variances, compared using the F-test for equality of variance between non-isogenic and isogenic lines, were smaller in the isogenic pairs, with all but the astrocyte staining meeting the threshold for significance (Fig. 7a-e). This finding was expected, given the single founder line and precisely targeted mutations. Surprisingly, however, the within-line variance metrics were reduced in the isogenic lines compared to the non-isogenic lines for all metrics, with only the microglia staining variances not meeting threshold for statistical significance (Fig. 7f-j). This lower variation was also evident in the qualitatively similar architecture of the developing organoid cultures across the isogenic lines (Extended Data Fig. 10 compared to Extended Data Fig. 1).

### Mutations in familial ALS genes give rise to distinct NMJ phenotypes

Having found deficits in muscle contractions as an initial ALS disease modeling phenotype, we focused our studies with the isogenic pairs on understanding potential structural contributions to explain the physiological impairment. We labeled neurons in isogenic organoid cultures by transduction with an *hSYN*::EGFP AAV9 vector (Fig. 8a) at week two and tracked neuronal growth longitudinally from weeks three to nine of culture. There was a significant growth of neurite area over time (*n*=6 organoid cultures obtained from 3 separate differentiations; linear model F=8.743, P=1.3×10^-10^) with no significant interaction between time and line (F=0.562, P=0.985), indicating similar rates of neurite growth between each pair of isogenic lines (Fig. 8b and Extended Data Fig. 11).

Because the ALS lines showed neither a deficit in neurite outgrowth nor a difference in percentage of skeletal muscle area (Fig. 6f) when compared to their isogenic controls, we examined the NMJs by transducing separate cultures with *hSYN*::EGFP at 13 weeks and labeling with α-bungarotoxin two weeks later. The number of labeled cells was similar among lines (*n*=6 organoid cultures obtained from 3 separate differentiations; Kruskal-Wallis, P=0.522; Extended Data Fig. 12a). The percentage of innervated NMJs, as defined by the number of EGFP+ and α-bungarotoxin+ puncta divided by the total α-bungarotoxin+ puncta, was reduced in the SOD1 and PFN1 mutant lines but not the TDP-43 mutant line (*n*=6 organoid cultures obtained from 3 separate differentiations; one-way ANOVA: F=5.938, P=6.3×10^-4^; two-tailed t-tests; TDP-43 pair: P=0.560; SOD1 pair: P=0.010; PFN1 pair: P=0.047; Fig. 8c). The size of the innervated NMJs, but not that of the uninnervated α-bungarotoxin clusters (Extended Data Fig. 12b), was reduced in the TDP-43 mutant line compared to its control, whereas the SOD1 and PFN1 pairs did not show effects of innervation state on NMJ area (*n*=6 organoid cultures obtained from 3 separate differentiations; one-way ANOVA: F=3.001, P=0.026; two-tailed t-tests; TDP-43 pair: P=0.009; SOD1 pair: P=0.203; PFN1: P=0.659; Fig. 8d, e). Thus, these results show a structural deficit in ALS NMJs, albeit without the temporal and cellular resolution of neuronal outgrowth and NMJ formation necessary to distinguish developmental versus degenerative contributions.

## Discussion

Although both iPSC-derived neuronal cultures and organoids yield synchronized networks of neurons^54, 62^, the ability to capture and interrogate specific synapses may prove critical to successful translational applications. Using a fusion of free-floating sphere culture followed by plating and growth under adherent conditions, we established sensorimotor organoid cultures and derived the NMJ together with other neuronal and non-neuronal cells that play cell autonomous and non-cell autonomous roles in sensorimotor diseases. In modeling ALS, for which NMJ loss is an early and critical component, we found evidence of compromised NMJs for several genes that span the broad categories of familial ALS-causing gene functions, namely, proteostasis, RNA binding proteins, and axonal transport^63^.

The reproducibility of neural organoids presents a challenge of uncertain magnitude^64^. Although recent studies are encouraging at least for more selectively patterned organoids^65, 66^, large variation in the types and numbers of cells generated – both among-line but particularly poor within-line reproducibility – has been a cause for loss of confidence in organoid models^67^, a hindrance in the transition from bench to clinic^68^, and a major focus of research effort^69–71^. To demonstrate the robustness of the sensorimotor organoid model, we characterized cultures generated using a total of 11 stem cell lines, including from healthy controls and ALS subjects, as well as gene-edited lines harboring ALS mutations and isogenic controls. We consistently observed a broad range of cells from neuronal and mesodermal lineages as well as motor neuron-dependent muscle contractions using the five non-isogenic lines studied, although with considerable variability. The variation of both sphere makeup and derived cell types was dramatically reduced in the six isogenic lines compared to either the single parental line or the other lines we previously examined. Both the among-line and, strikingly, the within line variances were reduced (Fig. 7), leading us to hypothesize that the proximity to clonal selection within only a small number of passages may be responsible. Intuitively, accumulating variation during iPSC replication and passaging might well increase the variation in differentiated organoid types. An alternative hypothesis, that the reduced variation in the isogenic lines reflects an unidentified consistent effect of gene editing at the three independent sites, seems less likely. The possibility that limited passaging after clonal selection can reduce variance will have to be tested formally in dedicated experiments and other organoid models, both neural and non-neuronal, with large numbers of independent differentiations to increase precision in variance estimates. If validated, the ability to reduce within-line variance by five to tenfold may prove broadly useful in organoid modeling, albeit requiring additional confirmation that genetic alterations due to clonal selection have been minimal^72^. Whether there may be implications for more routine differentiations of specific cellular types from iPSCs also remains to be seen, but less patterned differentiation strategies may be more vulnerable to greater iPSC variation.

Human *in vitro* disease modeling of the NMJ may help construct mechanistic hypotheses for early pathogenesis in neuromuscular diseases as well as facilitate the identification and validation of mechanistic targets and personalized preclinical therapeutic candidates. For ALS in particular, a growing body of evidence supports the importance of axonal biology in the disease^3, 73^, and the NMJ has provided the earliest and among the most important pathological and functional disease readouts for both mechanistic understanding and therapeutic development^1, 74, 75^. A recent study generated NMJs by co-culturing iPSC-derived motor neurons with myoblasts made from a single control iPSC line using a custom microfluidic device^8^. Optogenetic stimulation of motor neurons harboring ALS-causing mutations in TDP-43 yielded impaired entrainment of the control-derived muscle compared to stimulation of control motor neurons. In our data, we observed a reduction in the rate of large spontaneous contraction in all three ALS lines, a component of which may reflect additional non-cell autonomous effects of either the disease muscle itself or other cell types, as discussed below.

Comparison of pairs of isogenic lines bearing familial ALS mutations showed defects at the level of the NMJ and no impairment in neuronal outgrowth, supporting the primacy of the NMJ across ALS variants. One caveat was that the *hSYN*::EGFP virus did not report specifically motor neuron outgrowth, although the high percentage of motor neurons in the culture should nonetheless have preserved sensitivity to large effects in motor neurons. While increased numbers of lines will be needed for confirmation, the distinct phenotypes observed – namely a reduced percentage of innervated NMJs in the *SOD1*^G85R^ and *PFN1*^G118V^ mutations and a decreased area of innervated NMJs in the *TARDBP*^G298S^ mutation – are similar to prior reports demonstrating that different ALS genetic variants can affect the NMJ in distinct ways^76–78^. Future studies will be necessary to identify the mechanistic connections between these mutations and the NMJ physiological and structural phenotypes.

The strategy we employed to generate neuromesodermal precursors and subsequent organoids was similar to that recently described by Martins et al. using control human embryonic stem cell lines and an iPSC line^13^. While that technique yielded single, three-dimensional free-floating organoids, our cultured organoid strategy produced cultures that enabled live-cell imaging and facilitated quantification of cell types in all lines, measurement of within- and among line variation, and consequently disease modeling based on comparing NMJs derived from control and disease iPSC lines.

Limitations of our study included the ambiguity of developmental versus degenerative contributions to the observed phenotypes. In the future, longitudinal experiments may help address this issue. For example, such time course experiments might show a period of NMJ similarity between control and ALS organoid cultures followed by phenotype onset, and therefore support a degenerative etiology; however, we would be cautious in concluding this without a better understanding of the timing and duration of neurogenesis within organoid cultures among lines. Separately, longitudinal changes in organoids may not reflect *in vivo* development, in part due to the cellular stresses exerted by organoid culture^79^. Thus, longitudinal studies showing an apparent developmental phenotype could simply reflect differential sensitivity to stressors during what would otherwise be a common developmental program. These caveats aside, studies dedicated to the temporal evolution of the ALS NMJ phenotype combined with concomitant monitoring of key progenitors may provide insight into the mechanisms affecting NMJ physiology, number, and size.

While the structural complexity of the model may help better approximate features of the human disease, elucidating how different mechanisms act and interact – in disease-driving or compensatory ways – may prove challenging. For example, our observed reduction in muscle contraction within ALS organoid cultures may seem at odds with motor neuron hyperexcitability^80^. However, progressive hyperexcitability may lead to depolarization block and reduced motor neuron firing, consistent with other studies showing reduction in motor neuron firing, and thus explain the reduction in muscle contractions observed in this report^81, 82^. In this interpretation, hyperexcitability and apparent hypoexcitability due to depolarization block are on a continuum as opposed to opposite extremes. Alternatively, hyperexcitability could itself contribute to NMJ degeneration and weakened physiological function through a range of potential different implicated mechanisms, including ER stress, dipeptide formation, and TDP-43 pathology^83–85^.

Diseases of the sensorimotor system bear large non-cell autonomous effects from many cell types. For spinal muscular atrophy and ALS, astrocytes, microglia, and muscle may all contribute to motor neuron injury^86–89^; in chronic pain, the effects of microglia show unexpected sex-dependent differences^90^. Given the extensive number of causal and modifier genes for ALS, Charcot-Marie Tooth neuropathies, and hereditary sensory and autonomic neuropathies (HSANs), one might anticipate that non-cell autonomous contributions vary among disease and even disease subgroup. While co-culture experiments have demonstrated some non-cell autonomous toxicities^91^, specification of non-cell autonomous phenotypes and pathology can require elaborate interactions among multiple diverse cell types, for example, microglia, astrocytes, and neurons^92^. Compared with efforts to culture several cell types together^93^ or use remaining multipotent cells to perform sequential differentiations^12^, the organoid approach may actually improve variance by avoiding separate batch effects that arise from the generation of each cell type independently. We did not address the integration of cell types beyond motor neurons and muscle at the NMJ, such as to what extent the astrocytes or microglia exerted non-cell autonomous effects or whether the vascular cells contributed broad vascularization and even capacity for blood brain barrier modeling as has been seen in co-culture studies^94^. Our study provided only limited molecular and transcriptional resolution of cell subtypes present at later points in the culture. Given the observed NMJ phenotypes, we now anticipate that additional single-cell sequencing as well as tissue-specific cell sorting or RiboTag labeling^95^ of individual cell classes will help evaluate the presence and maturation of distinct cell types, as well as identify within-batch cell type-specific transcriptomic ALS features and thus help improve understanding of non-cell autonomous effects for different disease-causing mutations and genetic backgrounds.

In addition to motor neurons, the model generated dorsal spinal cord derivatives. We found nociceptor-like cells arranged in ganglia, and a large percentage of these cells yielded robust TTX-resistant sodium currents and capsaicin-induced calcium flux, substantial improvements in both percentage of cells responding and amplitudes over prior iPSC, lineage reprogramming, and organoid studies^96–99^. Although we have not yet applied the model to diseases with preponderant sensory phenotypes, it may prove useful for aggressive inherited conditions such as HSANs or acquired ones such as painful diabetic neuropathy^100, 101^.

The generation of distinct neuronal subtypes, such as nociceptors and motor neurons, provides an opportunity to test hypotheses of both disease heterogeneity and specificity. Our results suggest that different ALS variants may affect the NMJ in different ways, and larger numbers of distinct gene variant lines will be necessary to confirm this result. One would expect that motor neuron diseases would generally spare sensory neurons while sensory disorders, like HSANs, would not affect motor neurons. These hypotheses can now be investigated in a within-batch experimental setup. We propose that the sensorimotor organoid model will yield value in modeling a broad range of sensory and neuromuscular diseases.

## Author Contributions

J.D.P, D.M.D., A-C.D., A.H, Y.S., and B.J.W designed the experiments and wrote the paper. All authors reviewed and edited the manuscript. J.D.P, A-C.D., D.M.D., Y.S., J.H, and V.C. performed experiments. D.M.D. analyzed and interpreted single-cell RNA-seq data together with the Harvard Chan Bioinformatics Core. E.B., D.M.D., and A.H quantified muscle contractions. J.D.P., D.M.D., and A.H. quantified micrographs and performed statistical analyses.

## Acknowledgments

This work was supported by NIH grants F32-NS114319 (A.H.), DP2-NS106664 and K08-NS082364, the New York Stem Cell Foundation, and Target ALS (B.J.W.). B.J.W is a New York Stem Cell – Robertson Investigator. Electron microscopy was performed in the Microscopy Core of the Center for Systems Biology/Program in Membrane Biology, which is partially supported by an Inflammatory Bowel Disease Grant DK043351 and a Boston Area Diabetes and Endocrinology Research Center (BADERC) Award DK057521. Single-cell RNA-seq encapsulation and cDNA library preparation was performed at the Single Cell Core at Harvard Medical School, Boston, MA, cDNA libraries were sequenced at the Center for Cancer Computational Biology at the Dana-Farber Cancer Institute, and quality control and data analysis was done in collaboration with the Harvard Chan Bioinformatics Core at the Harvard T. H. Chan School of Public Health, Boston MA. Generation of isogenic ALS lines was performed at Harvard Stem Cell Institute iPSC Core Facility. We thank members of the Lagier-Tourenne and Albers labs as well as J. Sanes, P. Arlotta, and K. Hochedlinger for helpful discussion, C. Marques for assistance with figures, and A. Cohen for constructs.

## Online Methods

All experiments were conducted under protocols approved by the Massachusetts General Hospital Institutional Review Board and Partners Institutional Biosafety Committee. All methods were performed in accordance with relevant guidelines and regulations.

### Human iPSCs

Human iPSC cells included the previously available non-isogenic control lines 11a^102^ and FA0000011 (https://nindsgenetics.org/target-als-project), two familial ALS lines harboring mutations in *C9orf72* (19f) and *FUS* (MGH5b)^80^, and sporadic ALS line FA0000012 (https://nindsgenetics.org/target-als-project). Newly generated isogenic lines included TDP-43^+/G298S^, SOD1^+/G85R^, and PFN1^+/G118V^, as well as individual isogenic controls, in the FA0000011 iPSC genetic background.

### Gene Editing of Isogenic iPSC pairs

Gene editing of isogenic iPSC pairs was performed in collaboration with the Harvard Stem Cell Institute iPSC Core Facility. Briefly, the FA0000011 control iPSC line was characterized for morphology and pluripotency at passage 13 (Supplementary Data 1) and transfected with pCas9GFP (pCas9GFP was a gift from Kiran Musunuru, Addgene plasmid # 44719; http://n2t.net/addgene:44719; RRID: Addgene_44719)^103^ and independent guide RNAs cloned into pSPgRNA (pSPgRNA was a gift from Charles Gersbach, Addgene plasmid # 47108; http://n2t.net/addgene:47108; RRID: Addgene_47108; see Supplementary Table 2 for list of guide RNAs used)^104^ using Lipofectamine 3000. GFP positive cells were sorted and seeded at low density until single cell-derived colonies were formed. Clones were picked and cultured in a 96-well plate format for analysis of target mutations by Sanger DNA sequencing. Clones with a single point mutation in one of the target gene alleles and no modifications to the opposing allele were selected as heterozygous mutants and clones without any detectable modifications in any of the alleles of the target gene were selected as paired controls.

### Whole Exome Sequencing of Isogenic Pairs

Library preparation and exome sequencing were done by Genewiz Inc. using 150bp paired-end reads on an Illumina HiSeq sequencer. More than 94 million total reads were sequenced per library and aligned to GRCh37 using Edico Genome’s DRAGEN pipeline. Comparison of copy number variants was performed across lines with vcf-isec (r953). Multiple component analysis and clustering were completed using the FactoMineR (v. 2.3) and pheatmap (v. 1.0.12) R packages in R (v. 3.5.0).

### Cell Culture and Cultured Organoid Differentiation

iPSCs and ES cells were thawed and cultured as previously described^54^. In brief, iPSCs and ES cells were grown in six-well plates coated with matrigel (Corning, 354277) in mTeSR1 media (Stem Cell Technologies, 05850) until reaching 90% confluence, at which point they were differentiated or passaged.

iPSC cells that reached 90% confluency were enzymatically dissociated with StemPro Accutase Cell Dissociation Reagent (Thermo Fisher Scientific, A1110501) for 10 minutes at 37⁰C (day 0). Dissociated single cells were resuspended in organoid differentiation media consisting of STEMdiff APEL2 Medium (Stem Cell Technologies, 05275) and 5.32% Protein Free

Hybridoma Medium II (Thermo Fisher Scientific, 12040077), supplemented with the GSK-3 Inhibitor IX (0.5 μM) (Santa Cruz Biotechnologies, sc-202634), FGF2 (10 ng/ml) (Thermo Fisher Scientific, 13256029), Forskolin (20 μM) (Santa Cruz Biotechnologies, sc-3562) and Y27632 (10 μM) (Abcam, ab120129) and counted using a Countess II FL Automated Cell Counter (Thermo Fisher Scientific, AMQAF1000). Cells were then seeded as a 3D non-adherent culture at a density of 300,000 cells/ml for each well of an AggreWell 800 plate (Stem Cell Technologies, 34815), which had first been treated with Anti-Adherence Rinsing Solution (Stem Cell Technologies, 07010).

One day after seeding (day 1), the media was supplemented with 30% by volume base media and 30 ng/ml FGF2, 60 μM Forskolin, and 1.5 μM GSK-3 Inhibitor IX. From day 2 to day 6, daily media changes were performed: spheres were collected and centrifuged at 200 rcf for 5 minutes, and the media was aspirated and replaced with base media and FGF2 (10 ng/ml), Forskolin (20 μM) and GSK-3 Inhibitor IX (0.5 μM). On day 7, the spheres were counted with a Cytation 5 Cell Imaging Reader (Biotek), resuspended and centrifuged at 200 rcf for 5 minutes and plated at a density of 600 spheres per 35 mm dish on plasticware coated with Matrigel (Corning, 354277). From day nine onwards, the media was replaced daily with DMEM with High Glucose (Thermo Fisher Scientific, 11965092) supplemented with 2% Horse Serum (Thermo Fisher Scientific, 16050130) and 1% Pen/Strep (Life Technologies, 15070-063) for up to three months.

### Immunocytochemistry

Organoid cultures were fixed for 20 minutes at room temperature with 4% (w/v) formaldehyde (Thermo Fisher Scientific, 28908) and rinsed gently three times with PBS (Thermo Fisher Scientific, 10010049). Cultures were then permeabilized with 2% Triton X-100 (Millipore Sigma, 9400) in PBS for 20 minutes, followed by incubation with primary antibodies (Supplementary Table 3) in the same 2% Triton X-100 solution overnight at 37⁰C. Primary antibodies were removed the following day, and cultures were washed three times with PBS, followed by overnight incubation at 37⁰C with Alexa Fluor secondary antibodies (ThermoFisher Scientific) at a 1:500 dilution. Organoids were then washed three more times with PBS and mounted with Prolong Diamond (Thermo Fisher Scientific, P36962) per manufacturer’s instructions. Cultures were imaged the following day with either a Cytation 5 Cell Imaging Reader (Biotek) or an Image X-Press Micro Confocal (Molecular Devices). For analysis of NMJs, α-bungarotoxin directly conjugated with Alexa Fluor 647 (ThermoFisher Scientific, B35450) was resuspended per manufacturer’s instructions, diluted at 1µg/ml in PBS and applied to fixed organoids for 2h at room temperature. Immunostaining during the suspension period of organoid development (day 2) was performed by first plating the spheres and following the protocol described above. Quantification of individual cell populations was done through analysis of the whole well in MetaXpress (Version 6.5.4.532, Molecular Devices).

Immunostaining for FACS sorted cells was performed similarly, but secondary antibodies were incubated for two hours at room temperature prior to mounting with Prolong Diamond (Thermo Fisher Scientific, P36962) and imaging in Cytation 5 Cell Imaging Reader (Biotek).

### FACS Sorting

Dissociation of the organoid culture was done at week four by incubating the culture in 0.25% Trypsin-EDTA (Thermo Fisher Scientific, 25200056) for 20 minutes at 37⁰C. The resulting cell suspension was dissociated mechanically by pipetting ten times, followed by centrifugation at 200 rcf for 5 minutes. Immunolabeling was performed as previously described ^105^. Briefly, the cell pellet was resuspended into 200 µl of Fc-block antibody solution (1:100 of Human Fc Receptor Binding Inhibitor, Purified, Thermo Fisher Scientific, 14-9161-73; in PBS with 2% FBS, GE Healthcare, SH30910.03HI) for 30 minutes at 4⁰C while protected from light. Directly conjugated antibodies (NCAM or IG control) were then diluted in PBS with 2%FBS. The cell suspension was split in a 9:1 ratio, and 10% was reserved as IG control. The antibody solutions were either added to 180 µl (NCAM) or 20 µl (IG) of the cell suspension and were incubated protected from light for 30 minutes at 4⁰C. Finally, the cells were washed twice by adding PBS with 2% FBS and centrifuging at 200 rcf for five minutes before sorting.

FACS sorting was performed as previously described^54^ with either a BD FACSAria II or a BD FACSAria Fusion. After sorting, cells were maintained in DMEM with High Glucose (Thermo Fisher Scientific, 11965092) supplemented with 2% Horse Serum (Thermo Fisher Scientific, 16050130) and Y27632 (10 μM) (Abcam, ab120129) for one hour before fixing with 4% (w/v) formaldehyde (Thermo Fisher Scientific, 28908) and immunostaining.

### Analysis of Sphere Potency

Whole well acquisition was performed for spheres stained for TBXT, SOX2, and DAPI at day 2 using an ImageXpress Micro Confocal (Molecular Devices). MATLAB (version 2018b; MathWorks, Natick, MA) was used to stitch the individually acquired images, identify individual spheres, and mask the nuclear, SOX2, and TBXT channels. The nuclear area of each sphere and the percentage of each sphere covered by the SOX2 and TBXT masks were then calculated.

Potency of individual spheres was calculated as the difference between the percentages of SOX2 and TBXT mask areas for all spheres with less than 25% non-stained area, thus yielding a range from -100 (TBXT-only sphere) to 100 (Sox2-only sphere), with spheres expressing equal amounts of SOX2 and TBXT (neuromesodermal) having a score of 0.

### Single-cell RNA-seq

Single-cell RNA-seq was performed by microfluidic inDrop encapsulation, barcoding, and library preparation, as previously described^17, 106^ by the Single Cell Core at Harvard Medical School, Boston, MA. In brief, three independent biological replicates of the 11a iPSC line were differentiated for 16 days. Cells were then dissociated with Trypsin-EDTA and encapsulated into droplets using a microfluidic device. Each of the biological triplicates resulted in three 3,000 cell libraries, totaling 9,000 cells per biological sample. Next generation sequencing of the cDNA libraries was performed in collaboration with the Center for Cancer Computational Biology at the Dana-Farber Cancer Institute using an Illumina NextSeq 500 next generation sequencer.

### Analysis of Single-cell RNAseq Data

Bioinformatic analysis of single-cell RNAseq data was performed in collaboration with the Harvard Chan Bioinformatics Core at the Harvard T. H. Chan School of Public Health, Boston MA. Raw sequence reads were processed using the bcbio-nextgen single-cell RNA-seq pipeline (https://bcbio-nextgen.readthedocs.io/en/latest/contents/pipelines.html#single-cell-rna-seq). The pipeline uses tools from the umis repository (https://github.com/vals/umis) to generate a cell by gene count matrix. FASTQ files were formatted to parse out non-biological segments of the reads (i.e. cellular barcode, sample barcodes, and UMIs). Excess cellular barcodes were removed to reduce artifacts. The reads were aligned to the GRCh38 Ensembl Release 90 transcriptome with RapMap^107^. Duplicate UMIs were collapsed, and the number of reads per transcript were counted for each cellular barcode. Samples were assessed for quality and filtered using the distributions of reads per cell, UMIs per cell, genes per cell, mitochondrial ratios per cell, UMIs vs. genes detected, UMIs vs. read counts, and novelty scores.

Cell clustering was performed using the Seurat R package (v 2.3.4)^18, 108^ in R (v. 3.5.0). Cells with less than 200 unique genes were removed from the analysis, and genes expressed in less than five cells were filtered out. Raw expression values were log normalized and each gene was scaled and centered after regression of contributions from batch, cell cycle phase, total number of reads, and number of mitochondrial genes. We then performed principle components analysis of genes with highly variable expression across all cells (*n=*1,882) to identify sets of genes that captured transcriptional variation. Using the top 15 principle components, cells were clustered with a resolution setting of 1.6, and clusters were visualized using tSNE dimensionality reduction. Marker genes for each cluster were identified as genes expressed in at least 50% of cells in a cluster and enriched relative to all non-cluster cells based on the negative binomial model.

SPRING analysis was performed using the webtool^30^: number of PCA dimensions chosen was 50, with gene filtering settings of a minimum count of 3, a variability percentile threshold of 80%, and 5 nearest neighbors. No minimum was applied for cell filtering and a minimum of 3 cells was used for gene filtering.

### QPCR

Global gene expression analysis of 10-week-old organoid cultures was performed by qPCR. RNA was extracted with Trizol and 2 µg of RNA were converted to cDNA using the High-Capacity cDNA Reverse Transcription Kit (Thermo Fisher Scientific, 4368814). qPCR was performed with iQ™ SYBR Green Supermix (Biorad, 1708882) in a Biorad CFX96 with the following program: 95⁰C for six minutes followed by 40 cycles of 95⁰C (30 seconds), 55⁰C (60 seconds), 72⁰C (90 seconds) and 95⁰C (60 seconds). Gene specific primer sequences can be found in Supplementary Table 4, with exception of the *CX3CR1* primer set (Qiagen, QT00203434).

### Electron Microscopy

Electron microscopy was performed in collaboration with the Program in Membrane Biology’s Microscopy Core at Massachusetts General Hospital. Organoid cultures, grown in 35 mm plates, were fixed with 2.5% glutaraldehyde in 0.1 M sodium cacodylate buffer (pH 7.4, Electron Microscopy Sciences, Hatfield, PA) at least 1hr at room temperature on a gentle rotator. Fresh fixative was added, and specimens were allowed to infiltrate overnight at 4⁰C. The cells were rinsed several times in cacodylate buffer & target (contracting) regions identified & excised for analysis. While the entire cell sheet was processed as described below, priority for sectioning & analysis was placed on the targeted excised regions. Specimens were post-fixed in 1.0% osmium tetroxide in cacodylate buffer for 1 hour at room temperature and rinsed several times in cacodylate buffer. Samples were then dehydrated through a graded series of ethanol to 100%, dehydrated briefly in 100% propylene oxide, then allowed to pre-infiltrate overnight at room temperature in a 1:1 mix of propylene oxide and Eponate resin (Ted Pella, Redding, CA) on a gentle rotator. The following day, specimens were allowed to infiltrate several hours in fresh 100% Eponate resin. The targeted regions were placed, lying flat, into the capped ends of BEEM capsules, fresh 100% Eponate resin was added, and specimens allowed to polymerize 24-48 hours at 60⁰C. Remaining (non-priority) cell sheet pieces were transferred to coated glass slides with additional 100% Eponate resin, covered with coated glass coverslips, weights applied, and also allowed to polymerize 24-48 hours at 60⁰C (the latter specimen preparations were archived for future use). Thin (70nm) sections were cut using a Leica EM UC7 ultramicrotome, collected onto formvar-coated grids, stained with uranyl acetate and Reynold’s lead citrate and examined in a JEOL JEM 1011 transmission electron microscope at 80 kV. Images were collected using an AMT digital imaging system with proprietary image capture software (Advanced Microscopy Techniques, Danvers, MA).

### Pharmacological Analysis of Skeletal Muscle Contractions

Organoids at 8-9 weeks were imaged using phase contrast in a Cytation 5 reader (Biotek) with temperature and CO_2_ control (37⁰C, 5% CO_2_). Media was changed 1h prior to the experiment and a single site per well was used to establish a baseline of contractions immediately prior to compound addition. Botulinum neurotoxin type A (6.7 nM final concentration) (List Biological Laboratories, 130A) and Tubocurarine (12.5 μM) (Tocris, 2820) were resuspended in water, per manufacturer’s instructions.

### Phenotypic Analysis of Skeletal Muscle Contractions

Three biological replicates of each iPSC line were differentiated into organoids and maintained for 7-8 weeks. Six low magnification (4x) fields of view containing visible muscle fibers in brightfield were selected for each culture and imaged for a period of five minutes each.

### Contraction Quantification

Quantification of contractions for the pharmacological inhibition of contractile activity were done by optic flow. Raw videos were converted to 8-bit format, and then Fiji ^58^ was used to apply a FFT2D band pass filter (5 pixels to 40 pixels, no stripe suppression, 5% direction tolerance, and autoscale after filtering). For quantification of pixel movement as a measurement of contractions, the Gaussian Window MSE Optic Flow plugin of Fiji was used with the following parameters: Sigma of 4.00 pixels, Maximal Distance of 7.00 pixels (to which pixel movement was normalized). The average intensity over the whole flow vector video was measured over time.

For optogenetic stimulation, neuronal activity in organoid cultures was induced using the blue shifted channelrhodopsin TsChR2 driven by the neuronal specific promoter of Synapsin (h*SYN1*)(a gift from Adam Cohen)^56^. Areas of the organoid with both neurons and striated muscle were stimulated with pulses of blue light (405 nm, 0.2 Hz, 300 ms pulse width), triggering muscle contractions. Trackmate (Fiji) was then used to track cell body movement throughout the video with the following settings: default calibration settings, default crop settings, Log Detector with parameters of 20 µm blob diameter, and a 0.1 threshold. No initial threshold was set (all spots were kept). A HyperStack Displayer View was selected. The tracking settings were of a simple LAP tracker, linking maximum distance of 20 µm, Gap-closing maximum distance of 15.0 µm and a gap-closing maximum frame gap of 5 frames.

For the comparison of ALS and control lines, custom MATLAB (version 2018b; MathWorks, Natick, MA) scripts were used to identify pixel intensity variation between frames using frame subtraction. The absolute value of pixel intensity variation was then averaged for the frame. Large changes in the average pixel intensity per frame marked putative sites of muscle contraction. Each potential contraction was then manually verified by a blinded observer to exclude air bubbles, noise, and artifacts. The validated contractions were then readily classified as small (encompassing less than half of field of view) or large (encompassing more than half of field of view). The total number of contractile sites, total number of contractions, and the number of large contractions were measured blindly to line and disease genotype.

### Calcium Imaging

Organoid cultures were loaded with Fluo4-AM by incubating at 37⁰C for 90 minutes and rinsed three times with PBS. Cells were then transferred to a sodium-based extracellular solution containing (in mM): 140 NaCl, 5 KCl, 2 CaCl2, 1 MgCl2, 10 D-Glucose, 10 Hepes, pH 7.4. The organoid cultures were then imaged using a Nikon TI-Eclipse microscope and an Andor Zyla sCMOS camera with a PE4000 Cool-LED light source. Exposure times were 40-60 ms and images were taken every 0.5 sec. Capsaicin (1 µM) was added for 30 seconds after a two-minute baseline imaging recording. Individual cells were selected with NIS Elements AR software (v4.51.01, Build 1146, 64bit, Nikon) and calcium responses were calculated and graphed in MATLAB (version 2018b; MathWorks, Natick, MA) as ΔF/F to estimate comparative fluorescence intensity.

### Whole Cell Electrophysiology

Cultures were washed twice with the same solution described for calcium imaging prior to recording to remove media and debris. Cells were visualized using an inverted Nikon TI-Eclipse microscope and recordings were made using an EPC-10 amplifier (HEKA) controlled by PatchMaster software (PatchMaster v2x90.2, HEKA). During recording, cultures were continually perfused with aCSF containing (in mM): 127 NaCl, 3 KCl, 2 CaCl_2_, 1 MgCl_2_, 1.3 NaH_2_PO_4_, 10 Glucose, 25 NaHCO_3_ and gassed with 5%CO_2_ in oxygen. Intracellular pipet solution contained (in mM): 140 KMeSO_4_, 10 NaCl, 1 CaCl_2_, 1 EGTA, 3 MgATP, 0.4 Na_2_GTP, 10 HEPES. TTX (300nM) was applied for at least 10 min prior to recording to allow complete penetration into the tissue. Voltage-activated currents were elicited by 100 ms duration voltage steps to 0 mV from a holding potential of -80 mV.

### Neurite Outgrowth Analysis

Organoid cultures were transduced with an *hSYN*::EGFP AAV9 viral vector (a gift from Bryan Roth (Addgene viral prep # 50465-AAV9; http://n2t.net/addgene:50465; RRID:Addgene_50465) at week 2 of culture. A selection of 20 independent sites arranged in a fixed grid, constituting 7.8% of the total area of individual organoid cultures, was imaged longitudinally and bi-weekly, from week three to week nine in culture. Images were acquired with an Image X-Press Micro Confocal (Molecular Devices), and each individual site acquired as a z-stack of five individual images separated by 1 µm. Neurite outgrowth quantification was performed in Fiji (ImageJ 1.52p)^58^. Maximum intensity projections of each individual stack were loaded into ImageJ. First, neurites were identified by applying a 2 sigma gaussian blur, then applying a rolling ball filter with a 5 pixel radius to remove large objects. Long, non-circular neurites were then identified using Minimum Error automatic thresholding and selecting objects greater than 100 pixels^2^ and circularity values less than 0.5. Second, cell bodies were identified by applying a 2 sigma gaussian blur, identifying objects using Otsu automatic thresholding, and selecting objects greater than 100 pixels^2^ and circularity values greater than 0.3. Additional non-neurite noise resulting from autofluorescence of remaining large spheres was identified by applying a 10 sigma gaussian blur and Triangle automatic threshold. Neurite signal overlapping with either cell bodies or sphere autofluorescence was subtracted to yield the full mask of neurites, which was refined further to large neurite tracks by selecting objects greater than 1000 pixels^2^ and with circularity less than 0.2. For each neurite mask, the total area of the mask was measured. Neurite area for each image was loaded into R (v3.5.0) and total neurite area within each well was quantified by summing the area of each neurite mask. Neurite area was normalized to the initial timepoint for each well.

### Analysis of Percentage of Innervation and Area of NMJs

Organoid cultures were transduced with an *hSYN*::EGFP AAV9 viral vector (a gift from Bryan Roth (Addgene viral prep # 50465-AAV9; http://n2t.net/addgene:50465; RRID:Addgene_50465) at week 13 of culture. At 15 weeks of culture, organoid cultures were fixed with 4% (w/v) formaldehyde (Thermo Fisher Scientific, 28908) and rinsed gently three times with PBS. Organoid cultures were then labeled with a 1µg/ml solution of α-bungarotoxin directly conjugated with an Alexafluor^TM^ 647 dye (Thermo Fisher Scientific, B35450) in PBS, for 2h at room temperature. After incubation, excess dye was rinsed with PBS and the organoid cultures were imaged.

A random selection of 40 independent sites arranged in a fixed grid and constituting 15.6% of the total area of individual organoid culture wells were imaged for each organoid culture. Images were acquired with an Image X-Press Micro Confocal (Molecular Devices), and each individual site was acquired as a z-stack of five individual images separated by 1 µm. Quantification of EGFP expressing cells was performed manually, blinded to iPSC line and genotype. Detection of individual clusters of α-bungarotoxin, quantification of the intensity of α-bungarotoxin and EGFP, and quantification of the area of the α-bungarotoxin clusters were done with MetaXpress (Version 6.5.4.532, Molecular Devices). Analysis results were exported to R (v3.5.0). Clusters consistent with dimensions (10-50 µm) within the range of that of a human NMJ^109^ intensity of α-bungarotoxin labeling above the intensity threshold of 1000 were quantified for the total amount of α-bungarotoxin, with EGFP labeling above the intensity threshold of 5000 were considered innervated. Percentage of innervated NMJs was calculated as ratio between innervated and total α-bungarotoxin clusters per organoid culture. The areas of NMJs and uninervated α-bungarotoxin clusters were calculated as the average area of either innervated or uninnervated α-bungarotoxin clusters per organoid culture. Displayed data were normalized to the mean of matching isogenic controls.

### Statistics

All experiments and analyses were performed blind to cell line/drug identity. Sample sizes were chosen based on typical numbers in the field, with at least two independent differentiations for each experiment, with the exception of FACS sorting of neuronal cells. For all data, the Shapiro– Wilk test was used to confirm or reject normality. For normally distributed data, significance was assessed by one-way ANOVA and post-hoc two-tailed t-tests for pairwise comparisons. For non-normal data, the non-parametric Kruskal-Wallis test was used, with post-hoc Mann-Whitney tests.

To compare amplitudes of TTX-R sodium and potassium currents from individual cells grouped by the presence or absence of TTX-R currents, we performed Mann-Whitney tests and reported median and interquartile range (I.Q.R.). In Fig. 3h, group sizes were 4 and 10; U=0; P =0.002 for sodium currents and group sizes were 4 and 10; U=19; P =0.945 for potassium currents. For Fig. 5f, ANOVA (type 3 sum of squares in R) was used to compare contractile sites and contraction rates between control and ALS organoid cultures. For all statistics, significance was determined by 2-sided p-values under 0.05.

For the analysis of variance in non-isogenic and isogenic iPSC lines, measurements of individual sphere potency and area, as well as GFAP+ astrocytes, IBA+ primitive microglia, and SAA+ skeletal muscle area from mature organoid cultures were collected from 5-6 wells across 3 differentiation batches. For quantification of among-line variance, values for each well were normalized as fold change from the batch average across all lines. The mean value for each line was calculated and the variance of values among the 5 non-isogenic lines and 6 isogenic lines were compared using an F-test for Equality of Variances. For quantification of within-line variance, values for each well were normalized as fold change from the batch average for each line independently. The variance of values, that is the sum of squared deviations from the normalized average, were calculated for each line and the difference between average within-line variance among isogenic and non-isogenic lines was assessed using a one-way ANOVA.

### Data and Code Availability

Datasets, cell lines, and analysis tools will be available to the full authorized extent by request.

